# *spotter*: A single-nucleotide resolution stochastic simulation model of supercoiling-mediated transcription and translation in prokaryotes

**DOI:** 10.1101/2023.04.21.537861

**Authors:** William C. Hacker, Adrian H. Elcock

## Abstract

Stochastic simulation models have played an important role in efforts to understand the mechanistic basis of prokaryotic transcription and translation. Despite the fundamental linkage of these processes in bacterial cells, however, most simulation models have been limited to representations of either transcription *or* translation. In addition, the available simulation models typically either attempt to recapitulate data from single-molecule experiments without considering cellular-scale high-throughput sequencing data or, conversely, seek to reproduce cellular-scale data without paying close attention to many of the mechanistic details. To address these limitations, we here present *spotter* (Simulation of Prokaryotic Operon Transcription & Translation Elongation Reactions), a flexible, user-friendly simulation model that offers highly-detailed combined representations of prokaryotic transcription, translation, and DNA supercoiling. In incorporating nascent transcript and ribosomal profiling sequencing data, *spotter* provides a critical bridge between data collected in single-molecule experiments and data collected at the cellular scale. Importantly, in addition to rapidly generating output that can be aggregated for comparison with next-generation sequencing and proteomics data, *spotter* produces residue-level positional information that can be used to visualize individual simulation trajectories in detail. We anticipate that *spotter* will be a useful tool in exploring the interplay of processes that are crucially linked in prokaryotes.

## INTRODUCTION

Recent work from a number of groups has demonstrated that stochastic simulation methods can be used effectively to model bacterial transcription and translation. While earlier models established the power of stochastic methods in large systems at relatively coarse resolution^1,2^, recent studies have focused on the step-by-step progress of RNAPs and ribosomes along their respective templates. Sophisticated models of transcription have incorporated experimentally-observed phenomena, including pausing and backtracking, in attempts to develop physically-plausible, sequence-dependent rates of translocation that are consistent with data obtained from single-molecule studies^3–4^ or to represent RNAPs in traffic^5–8^. Several models featuring the interplay between transcription and supercoiling have emerged as well^9–14^, reflecting recent experimental evidence that DNA supercoiling can affect both transcription initiation^15^ and elongation rates^16^. Separately, powerful machinery for the simulation of translation has also been developed, in which codon-specific ribosomal dwell times are assigned based on either tRNA availability^17–20^ or ribosomal profiling data^21–22^. Many translation models include an impressive range of reactions, accounting explicitly for elongation factors^23^ or modeling collision-stimulated termination^24^. Finally, while most studies have considered transcription *or* translation separately, some detailed simulation models focused on transcription^25^ have been extended to include translation^26^. Perhaps most interestingly, systems in which co-transcriptional translation can occur have also been reported^27–29^; such models are important because transcription and translation in bacteria occur in the same cellular compartment, meaning that bacterial ribosomes have access to mRNA even as it is being transcribed.

Despite these advances, to our knowledge, no simulation model that can simultaneously represent transcription, DNA supercoiling, and translation—processes that are all fundamentally interlinked in the bacterial cell—is currently available. Moreover, those simulation models that do handle a subset of these processes are often not available in user-friendly, flexible implementations. To address these limitations, we describe here *spotter*, an integrated stochastic simulation model of prokaryotic transcription and translation that provides a ready-to-use toolkit. *spotter* employs a new model of sequence-dependent transcription that allows nucleic acid hybridization energies to be combined with experimental sequencing data and includes sophisticated models for RNAP pausing, backtracking, and rotational relaxation. In addition, *spotter* provides a detailed representation of DNA supercoiling (including its modulation by the action of processive topoisomerases), and a treatment of translation that is informed by both tRNA competition and ribosomal profiling data (Figure 1; Figure S1). Crucially, because all of these processes contribute to a single, integrated model, *spotter* is uniquely positioned to explore systems that cannot be treated in currently-available software, allowing, for example, investigation of the relationship between DNA supercoiling and RNAP backtracking and the modulation of RNAP elongation rates by co-transcriptional translation.

**Figure 1.**
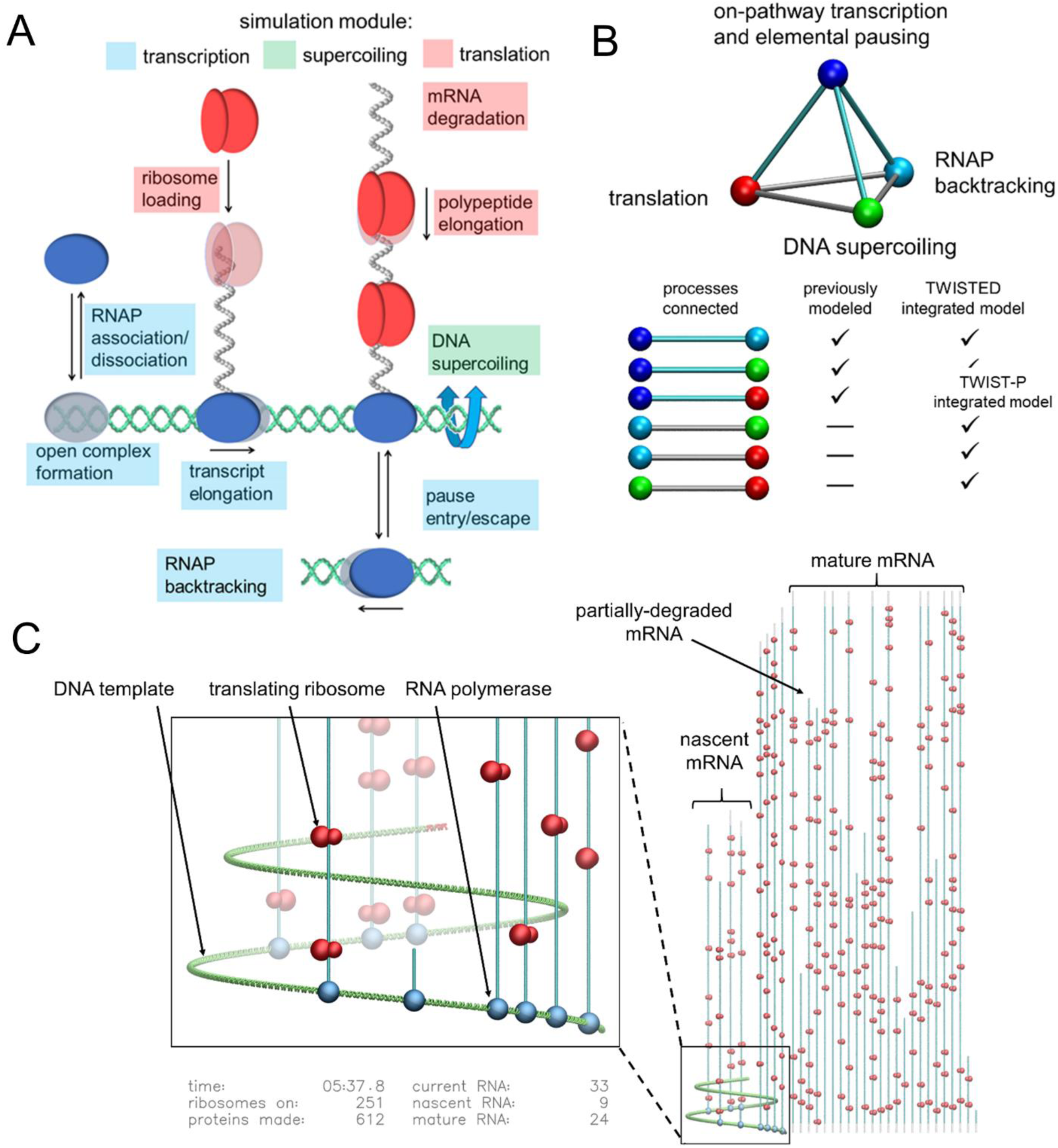
Overview of *spotter*. **A.** Core processes represented in the integrated model. Essential processes occurring on DNA and RNA can be grouped conceptually into “modules”: a “transcription module” (processes boxed in blue) includes reactions representing RNAPs’ association with and progress along the DNA template, as well as associated RNA production; a “supercoiling module” (boxed in green) accounts for events that change the local supercoiling density of the DNA template; and a “translation module” (boxed in red) includes reactions representing the production of polypeptides, which can occur co-transcriptionally on template mRNA in simulations, as well as RNA degradation. **B.** Novel connections between processes modeled in *spotter* simulations. Tetrahedron edges in cyan represent processes (tetrahedron vertices) that have been modeled simultaneously in pairwise fashion in previous simulation work. All processes can be modeled in a single simulation with the *spotter* model; newly-connected processes are represented by tetrahedron edges in gray. **C**. Representative snapshot from a simulation of the *E. coli alaS* transcription unit. Images were produced with the *spotter* visualization toolkit from a simulation of a single copy of the *alaS* transcription unit set to produce mRNA and protein at physiological levels. In the image at right, template DNA (green helix) is transcribed by RNAPs (blue spheres). The resulting mRNA (gray, with the *alaS* gene overlaid in cyan) is oriented so that 5’ to 3’ movement of translating ribosomes (red) occurs vertically down the page. The complete set of mature mRNAs appears to the right of the DNA template; as is indicated in the close-up at left, both translation and degradation of mRNA can occur co-transcriptionally in simulations. The visualization kit also displays continuously-updated RNA and protein inventories during each simulation trajectory (example at bottom left

This manuscript is structured as follows. In Methods we provide a description of *spotter*’s overall structure, and its conceptual separation into dedicated modules describing transcription, DNA supercoiling, and translation. Methods provides concise descriptions of how input data for simulations might be constructed, but the full details of these issues are deferred to the Supplementary Material. In Results, we describe a series of simulation test cases, each of increasing complexity, that serve to illustrate *spotter*’s capabilities; we build from simulations of transcription by lone RNAPs to simulations of coupled transcription and translation by multiple RNAPs on polycistronic operons. In Discussion, we seek to place *spotter*’s capabilities in context, revisiting the examples described in the Results and identifying both those aspects of its programming that have been borrowed from earlier models, and those aspects that, to our knowledge, are fundamentally new.

## METHODS

The following sections present the *spotter* simulation model in some detail, but a complete account of the model—including the rationale behind each aspect of its design, the detailed equations used to determine instantaneous reaction rates, and comparisons with other simulation models—is presented in the Supplementary Material. Complete instructions for generating the files required for a *spotter* run, choosing among available model options, running simulations, and visualizing output trajectories are provided in documentation accompanying the *spotter* source code – all of which is written in the C programming language – and can be found at the following GitHub repository (https://github.com/Elcock-Lab/spotter).

### General features of the *spotter* simulation model

*spotter* simulations are carried out at the level of the transcription unit or operon and include reactions contributing to three fundamental phenomena: (a) the generation of mRNAs from a DNA template by RNAP-mediated transcription, (b) the modulation of DNA supercoiling by transcription events, and by the action of topoisomerases, and (c) the translation of mRNAs into proteins by ribosome-mediated translation. All the elementary reactions that contribute to these phenomena are simulated stochastically using the Next Reaction Method of Gibson and Bruck^30^ as developed from the Gillespie algorithm^31^. These reactions are organized around two major “object” types, the DNA template and the mRNA template, each of which is represented in the simulation model as a one-dimensional lattice. The DNA template is present in a single copy of invariant length and is subdivided into topologically-insulated domains whose boundaries are determined by the position of transcribing RNAPs and/or by fixed domain boundaries specified by the user. The RNA template, in contrast, can be present in multiple independent copies with lengths dependent on the progress of: (a) the RNAP that creates them, and (b) the degradation machinery that destroys them. In the case of polycistronic operons, each mRNA template is further divided into regions corresponding to each gene to enable the production of different types of protein from the same RNA; there is no limit on the number of genes that may be contained within a single mRNA template, so even the most complex bacterial operons can be modeled.

*spotter* simulations describe the actions of another set of “objects” that bind to (DNA and RNA), translocate along (DNA and RNA), degrade (RNA), rotate around (DNA), or enzymatically relax (DNA) the templates with which they interact. A critical rule in the simulations is that these DNA- and RNA-binding objects cannot occupy positions already rendered physically unavailable by the binding of another object. To enforce this rule, each object is assigned a characteristic footprint: RNAP, topoisomerase I, and gyrase, for example, are assigned footprints of 35, 50, and 150 bp of DNA, respectively, and the ribosome is assigned a footprint of 30 nt of mRNA; all of these estimates are based on experimental data^32–36^ but they can be changed by the user.

A variety of files and parameters serve as input to the simulations. The rates of transcription and translation elongation steps are determined by dwell-times specified by the user for each position along the gene or transcript, respectively. At the simplest level, a single value can be used for all positions as input in order to provide completely flat transcriptional or translational landscapes; alternatively, completely arbitrary sequences of values can be input. In most cases, however, we anticipate that the user will be most interested in inputting dwell times that accurately represent those of known genes or transcripts. To this end, *spotter* provides preprocessing utilities that can derive dwell times for both transcription and translation reactions from experimental NGS data. Rates for transcription reactions, for example, can be generated from user-supplied files that represent the DNA sequence and available experimental NET-seq data to match a mean desired rate of unobstructed transcriptional elongation. Rates for translation reactions, on the other hand, can be derived from a user-supplied RNA sequence together with ribosomal profiling (ribo-seq) data to match a mean desired rate of unobstructed translational elongation. Default rate constants are provided for all other reactions, but these are also modifiable in the input files provided to the program. With all rate constants specified, the user only needs to provide a command file that specifies the desired duration of the simulation, the frequency with which data are saved to file, etc. We note that the present version of *spotter* is designed only for simulations at 37° C.

*spotter* produces a wide variety of outputs. These include inventories and positions of mRNAs, proteins, and DNA-bound RNAPs at user-selected time-points; in addition, individual trajectories can be used to generate “sequencing” data, which can be collated from sets of trajectories using a series of utility programs for the processing of simulation output, dwell-time probability distributions, and kymographs (as in Figures 5-9). Finally, *spotter* provides routines for making movies from simulation trajectories using the widely-used molecular graphics program VMD.^37^ In particular, users can choose to make “system-level” movies that represent the entire transcription unit, all of its extant RNAs, and all active ribosomes (examples include Movies S1-3) or “zoomed-in” movies that instead represent only a portion of the transcription unit and omit the RNA and ribosomes in order to display changing supercoiling densities, RNAP pause states, and RNAP rotation. Documentation describing the commands necessary to prepare simulation input files, run simulations, process simulation data, and generate movies from trajectories is included with the *spotter* source code.

In what follows, we outline each of *spotter*’s three main modules and describe how they might be used. Since the example applications described in Results all involve *spotter* simulations of *E. coli* genes, we emphasize how experimental data already available for *E. coli* can be used to set the input parameters that are provided to each of *spotter*’s modules. But, since *spotter* also allows users significant leeway in allowing both real and artificial systems to be simulated, further examples of how the input parameters can be specified – independent of data from *E. coli* – are provided in the Supplementary Material and in the code documentation.

### Transcription module

#### Overview

In developing *spotter* we focused on building in a flexible model of transcription that includes not only conventional on-pathway events but also those off-pathway events that allow a RNAP to enter a variety of paused or backtracked states. Importantly, many of the reactions that are handled by the transcription module are directly affected by, and affect, those occurring in a separate module that explicitly monitors the state of supercoiling of the DNA template (see below). The full set of reactions and states available to elongating RNAPs in *spotter*’s transcription module are illustrated in Figure 2; the initiation and termination stages of transcription simulations are discussed in detail in Supplementary Material. Briefly, an RNAP engaged in on-pathway transcriptional elongation advances along its DNA template via a reaction that moves it from a pre-translocated state to a (nucleotide-free) post-translocated state positioned one base-pair downstream (yellow arrow in Figure 2). It then enters a pre-translocated state at this new position through a consolidated reaction that describes both nucleotide addition and catalysis (green arrow in Figure 2). This cycle repeats for as long as the RNAP remains on-pathway. The cycle can be interrupted, however, if the RNAP instead enters a so-called “elemental pause”^38^ (orange arrow) in a reaction that competes with the forward translocation reaction and takes the RNAP into an “off-pathway” state that is maintained until it undergoes an exit reaction (blue arrow) that returns it to a pre-translocated state.

**Figure 2.**
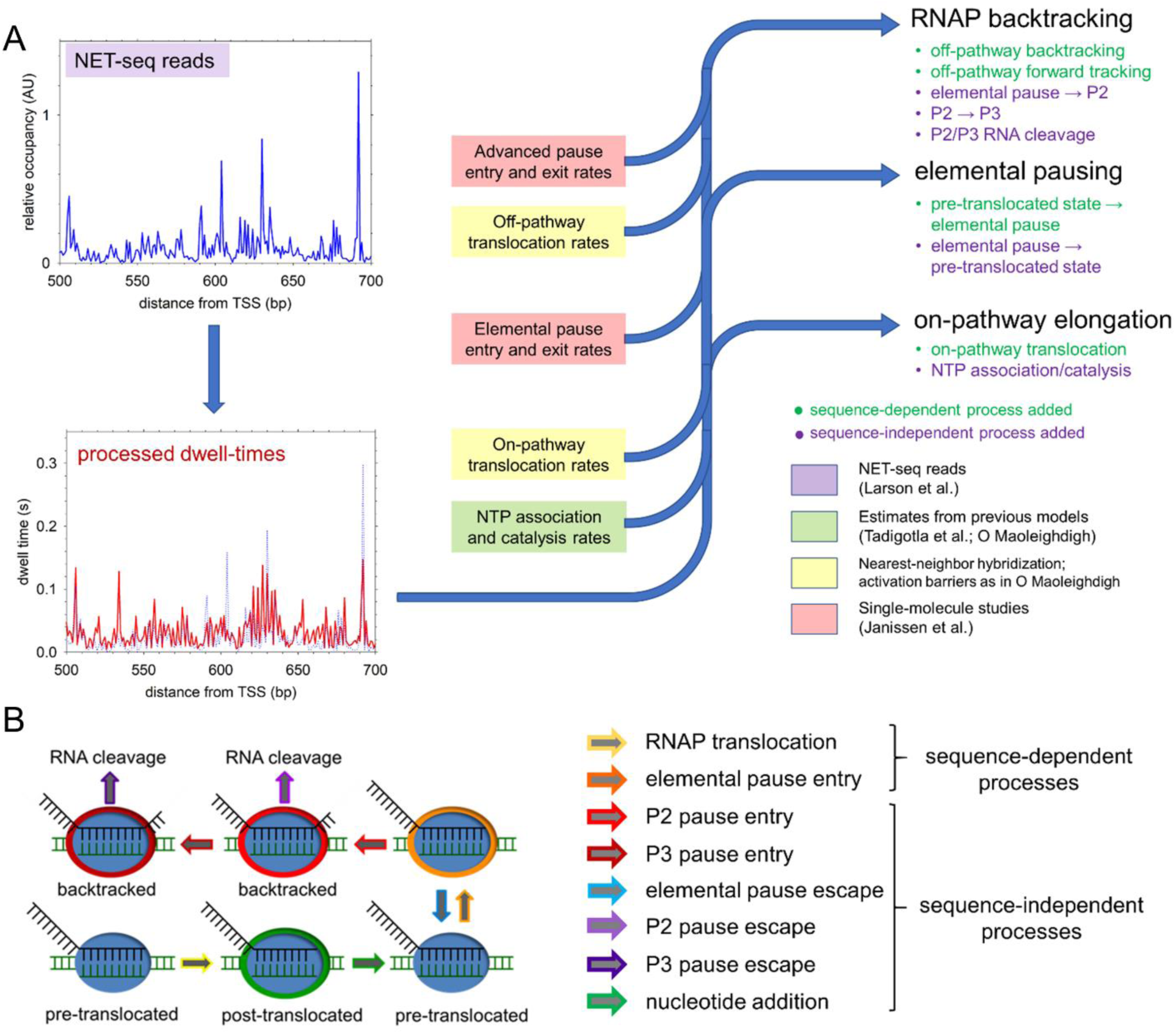
Features of the *spotter* transcription module. **A.** Incorporation of previous experimental and theoretical work in the multistate transcriptional model. All simulation models begin with NET-seq data from Larson et al. (GEO accession GSE 56720); plotted in blue at upper left); NET-seq reads are processed to determine sequence-specific dwell times (red line at lower left) at all positions in the genome of interest as described in Methods. The flow chart at right indicates the data incorporated at each stage in the development of the transcriptional model. **B.** Schematic of the multistate model. On-pathway RNAPs (blue ovals) may be pre-translocated (no outline) or post-translocated (green outline); in the latter state an elemental pause entry reaction competes with forward translocation. Advanced pause states include P2 (indicated in red outline), accessed from the elemental pause state by a P2 entry reaction (red arrow), and P3 (indicated in dark red outline), accessed from the P2 state by a P3 entry reaction (dark red arrow). RNAPs in the P2 or P3 state can back- or forward-track at all positions provided that they do not advance beyond the position in which they first entered an elemental pause. In simulations, RNAPs in state P2 or P3 return to on-pathway transcription (in the pre-translocated state) through RNA cleavage reactions^30^ that remove backtracked RNAPs’ 3’ overhangs (see Figure S2 for a schematic of available backtracking reactions).

While the ubiquitous elemental pause state identified experimentally is not long-lived (< 1 s), it is considered a necessary precursor to longer-lived pause states^38^ that are also included in the model. Access to these pause states is depicted schematically in Figure 2: an RNAP in the elementally-paused state first progresses to a longer-lived pause state P2 (outlined in bright red), from which it can in turn gain access to an even longer-lived pause state P3 (outlined in dark red). RNAPs in these advanced paused states are also subject to off-pathway back-tracking and forward-tracking translocation reactions (see below). Escape from advanced pause states is thought to require cleavage of extruded RNA by the polymerase itself or (more efficiently) by the enzymes greA and greB^39,40^ We model this escape process as a consolidated “exit” reaction in which RNA cleavage and the return of the RNAP to on-pathway transcription occur simultaneously. Exit from P2 or P3 can occur at any of the sites to which an RNAP has backtracked, where, upon RNA cleavage, it enters the pre-translocated state at that position (Figure S2).

There are several ways in which rates for each of the above reactions can be assigned by the user (see Supplementary Material), but the essential input provided by the user to the transcription module is the mean dwell time spent by an RNAP at each position along the DNA template. *spotter* then, by construction, ensures that these dwell times are reproduced exactly by stochastic simulations that include only a lone RNAP transcribing in the absence of any topological barriers that introduce supercoiling effects. This feature makes *spotter*’s transcription module well suited to producing transcription trajectories that explicitly reproduce experimental RNAP occupancy data. As is shown in Results, however, supercoiling-related effects and/or the interplay of multiple RNAPs transcribing the same transcript can cause the effective dwell times to deviate from their input values.

#### Use of NET-seq data to determine relative RNAP dwell times

Given that our original interest in developing *spotter* was to model transcription and translation in *E. coli*, all of the simulations reported here use sequence-specific dwell times that have been derived from analysis of NET-seq data reported by the Greenleaf, Landick and Weissman groups^41^. NET-seq is an NGS-based experimental method that specifically identifies the 3’ end of nascent transcripts and so provides a read-out of those locations on a DNA template where RNAPs tend to spend most of their time. Following Larson et al, we normalize the NET-seq read counts for each gene of interest using boundary definitions taken from RegulonDB for *E. coli* strain MG1655^42^; this normalization accounts for differences in transcript abundance and isolates intragenic differences in relative dwell times at positions within the gene. We next use an energy function derived by Larson et al.^41^ from their complete set of gene-normalized dwell times in order to assign relative dwell times to all positions on the template of interest based on the sequence of a 16-bp window surrounding the RNAP active site at each position (see Supplementary Material). The processed dwell times resulting from the application of this energy function (Figure 2A, at left) are designed to capture variations in dwell time intrinsic to sequence rather than variations that may be specific to the genomic context.

#### Calculation of RNAP translocation rates

Following others^3,4^, *spotter* bases its calculations of RNAP’s translocation rates on the estimated relative thermodynamic stabilities of the pre- and post-translocated states. Specifically, for each position in the gene of interest, the energies of the pre- and post-translocated states are calculated using nearest-neighbor (NN) model thermodynamic data for DNA-DNA^43^ and RNA-DNA hybridization^44^ at 37 °C. As previously suggested^4,45^, energies are calculated assuming a 12-bp transcription bubble and a 9-nt RNA-DNA hybrid in the pre-translocated state (the bubble is depicted schematically in Figure 2B); as with almost all parameters input to *spotter*, however, the size of the transcription bubble can be adjusted by the user. During forward translocation, the length of the RNA-DNA hybrid is reduced by 1 bp in preparation for addition of the next nucleotide and this is accompanied by rewinding and unwinding of the DNA by 1 nt at the upstream and downstream edges of the bubble, respectively. As is detailed in the Supplementary Material, we follow previous work^4^ in modeling rates of translocation with a Kramers form with an energetic barrier assigned as the mean of the pre- and post-translocated energies plus a global constant of 5 k_B_T. We incorporate parameters from previous experimental and theoretical work (Figure 2A, blue arrows at right) aimed at characterizing on-pathway translocation and nucleotide addition rates.

In addition to on-pathway translocation, RNAPs in advanced paused states can also translocate off-pathway with so-called back-tracking and forward-tracking reactions (Figure 2B; Figure S2). Rates for these reactions are determined from the same DNA-DNA and DNA-RNA hybridization energies^43,44^ calculated above for the pre- and post-translocated sites following the method outlined in previously reported simulation models^4,5^.

#### Assignment of rates to RNAP pause-entry and pause-exit reactions

To prevent an explosion in the number of parameters required of the user, certain rates assigned in *spotter*’s transcriptional module are, in the current version of *spotter*, assumed to be sequence independent. These are the following rates: (1) k_e1_, the rate at which an RNAP returns from the elementally-paused state, EP, to the pre-translocated state on the main pathway; (2) k_p2_, the rate at which an RNAP moves from the elementally-paused state to the longer-lived pause state P2; (3) k_p3_, the rate at which an RNAP moves from state P2 to the still-longer-lived pause state P3; (4) k_e2_, the rate at which cleavage occurs on an RNAP in state P2, returning it to the position it has moved to during backtracking; and (5) k_e3_, the corresponding rate, but for exit from state P3. The assumed sequence-independence of these rates is based on single-molecule experiments conducted on *E. coli* RNAP by the Dekker group^39^; rates for all five reactions were taken directly from that study and are set to 1.77 s^-1^, 0.2 s^-1^, 0.022 s^-1^, 0.241 s, and 0.01 s^-1^, respectively. Each of these parameters remains freely adjustable by the user.

While many of the above rates are assumed to be position-independent, the central assumption that we make here is to adjust independently the rate, k_p_, at which a pretranslocated RNAP enters the elementally-paused state at each position in order to ensure that the mean dwell time at that position matches the NET-seq-derived dwell time. This is achieved using the decomposition of the net elongation rate into a sum of exponential terms outlined by Janissen et al.^39^; full details of these calculations are provided in the Supplementary Material. In the example applications reported here, we have found that the decomposition always results in a physically realizable value of k_p_: for the genes studied here, the maximum and minimum derived rates for k_p_ are ∼1300 and 0.0005 s^-1^, respectively.

### Supercoiling module

#### Overview

The description of DNA supercoiling included in *spotter* is intended to be sophisticated: it directly accounts for the known bi-directional interplay between supercoiling and transcription, and it explicitly incorporates the presence of topological insulators and the effects of topoisomerases. Figure 3 illustrates the full set of reactions handled by *spotter*’s supercoiling module. The template DNA can be partitioned into topologically-insulated domains, i.e. regions of the DNA template enclosed between topological insulators that act as barriers to the diffusion of supercoiling; insulators can include actively transcribing RNAPs and/or additional barriers that define the boundaries of a larger topological domain. During *spotter* simulations, the supercoiling density (σ) of a given domain is updated immediately upon the occurrence of any event that changes it. These events include: (a) the translocation of an RNAP along the DNA template, (b) the rotation of an RNAP around the DNA template, and (c) the activity of the enzymes topoisomerase I and gyrase, which act to relax supercoiled DNA (at start of blue arrows in Figure 3). An event that updates the supercoiling density within a domain will, as a consequence, also necessitate updating the net torque experienced by adjacent RNAPs which, in turn, will require updating the rates of the torque-sensitive reactions available to those RNAPs. The latter include rates of pause entry and on- and off-pathway translocation (outlined in the flowchart).

**Figure 3.**
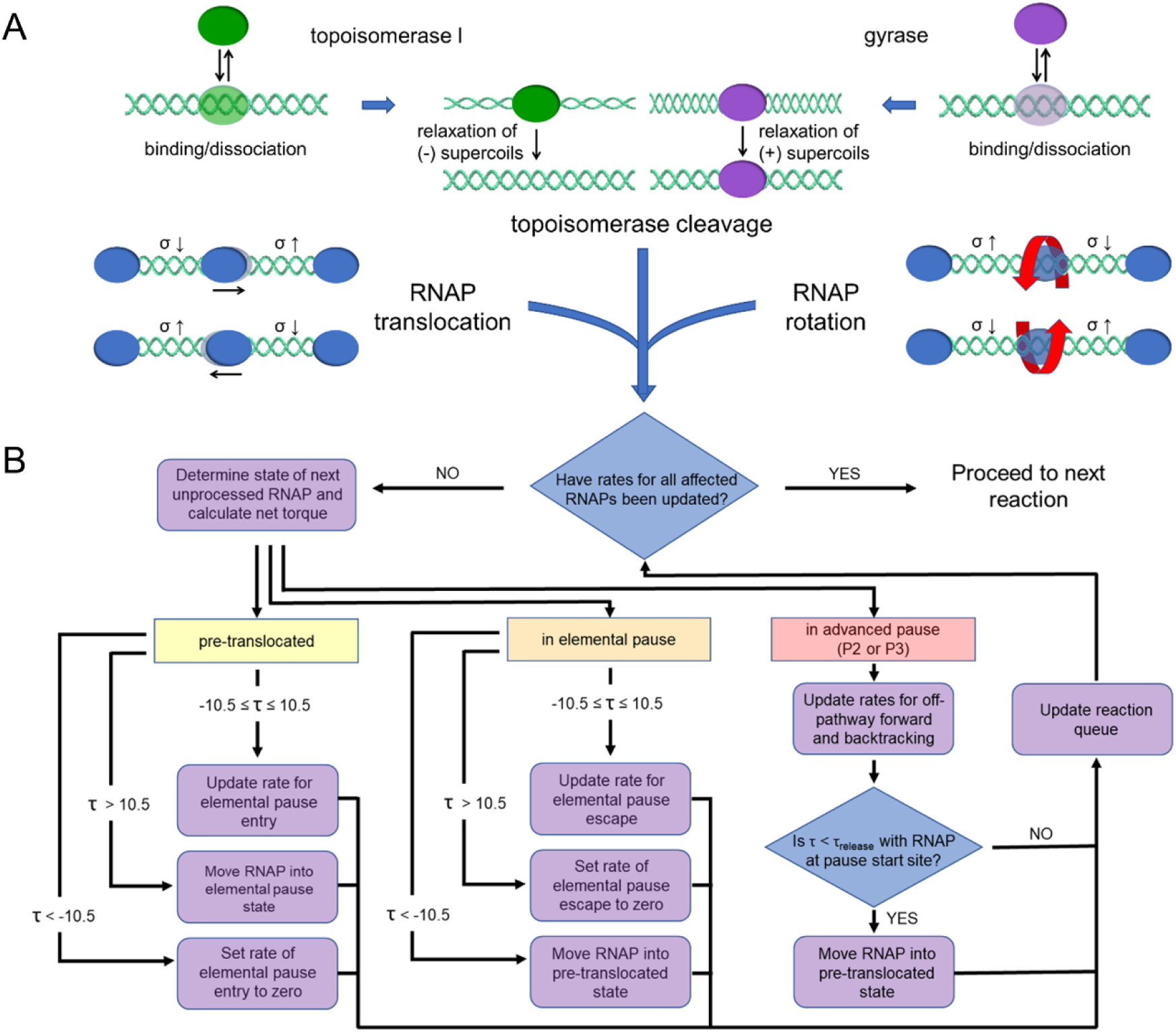
Central features included in the supercoiling module. **A**. Modulators of supercoiling. At the top of the figure (at blunt ends of convergent blue arrows), processes producing changes in local supercoiling density are depicted: these are (clockwise, from left) RNAP translocation, which decreases and increases σ in domains up- and downstream (respectively); topoisomerase activity, which increases (topoisomerase I) or decreases (gyrase) σ in the domain within which the enzyme acts; and RNAP rotational relaxation, which acts to equilibrate supercoiling density in the region within which an RNAP is located. **B.** Downstream effects of supercoiling modulation. The flowchart at the bottom of the figure illustrates the effect of changes in supercoiling density on reactions included in the transcription module. Because rates of nucleotide association and catalysis are assumed to be insensitive to supercoiling, RNAPs in the post-translocated state are unaffected by supercoiling. If an RNAP is in any other state (yellow, orange, and red boxes), the rates of reactions available to that RNAP are modified as indicated in the flowchart. We employ the fits determined by Heberling et al. to data from Wang et al., adjusted by NET-seq-derived dwell times as described in the main text and Methods in order to calculate new rates for all reactions. Reaction times are adjusted, created, or eliminated as indicated in the chart using previously-scheduled reaction times and new and updated rates according to the next-reaction method of Gibson and Bruck. The process of rate-adjustment is continued iteratively until rates for all RNAPs (up to three) affected by changes in σ have been adjusted.

#### Effects on transcriptional reactions

Following previous models of supercoiling’s effect on RNAP progress^9,10^ which were themselves derived from pioneering single-molecule studies of transcription under torsion^46^, a key variable that *spotter* tracks during simulations is the torque experienced by each transcribing RNAP. We calculate this torque from the difference of the supercoiling densities in the adjacent up- and downstream insulated domains, and we use previously-reported fits to single-molecule data^9^ to update both the probability of entering an elemental pause (by updating the pause entry rate, k_p_) and the duration of that pause (by updating the rate of pause escape, k_e1_) whenever the torque changes. The resulting reactions are then automatically rescheduled within *spotter*’s current roster of reaction events using the Next Reaction Method. Because, unlike previous transcriptional models that have included DNA supercoiling^9–14^, *spotter* includes position-specific elemental pause entry rates, we adjust the *relative* rate of pause entry at that position (see Supplementary Material).

Because *spotter* also appears to be unique among supercoiling simulation models in including the long-lived P2 and P3 pause states associated with RNAP backtracking, we also model the effect of σ differences in the domains up- and downstream of a backtracked RNAP on off-pathway rates of forward- and back-translocation. The adjustments of rates of elemental pause entry and exit reactions are handled as in the elemental-pause-only model (paths descending from yellow and orange boxes in the Figure 3 flowchart). In addition, however, supercoiling affects the rate at which RNAPs in state P2 or P3 back- and forward-track (paths descending from the red box in the flowchart at right): the zero-torque rates for these reactions are therefore adjusted according to a velocity-dependence curve (a fifth-order polynomial) previously used to calculate RNAP on-pathway translocation rates in a simulation model that did not include advanced pause states^9^ (Figure S7).

Finally, because recent single-molecule studies suggest that backtracked RNAPs in long-lived pause states do not resume active transcription simply by returning to their initial pause sites^39,40,46^, additional mechanisms for the supercoiling-mediated “rescue” of backtracked RNAPs appear to be necessary to prevent template congestion in physiologically-relevant conditions in which more than one RNAP is actively transcribing an operon. As is described in “Results” below, *spotter* also allows users to set a “release torque threshold” or minimum assistive torque that, when experienced by an RNAP in an advanced pause that has tracked back to the template position of its original pause, allows the RNAP to resume productive elongation from that position. The application of the release torque allows the cleavage-independent return to on-pathway transcription for RNAPs in the presence of other RNAPs transcribing the same template. Full details of the torque-dependence of all reactions are presented in the Supplementary Material.

#### RNAP rotational relaxation

The treatment of RNAPs as topological insulators in *spotter’s* model of DNA supercoiling reflects the very limited rotational freedom likely to be experienced by an RNAP with a lengthy nascent mRNA and substantial ribosomal cargo in tow^47^. *spotter* allows transcribing RNAPs to undergo relaxation reactions that rotate it around the DNA axis in the direction that equilibrates supercoiling density in the two domains on either side of it. As is described fully in the Supplementary Material, RNAP rotation is discretized so that each such reaction event results in the rotation by a user-specified angle; in all simulations reported here we used our default value of 3.43°, which is set so that 100 events are required to rotate by one turn of the DNA double helix. Rotation reaction rates are modeled as functions of: (a) the torque experienced by the RNAP as a result of differences in up- and downstream supercoiling densities, and (b) the viscous drag that the entire elongating complex is subject to, accounting for the RNAP itself, its RNA transcript, and the ribosomes translating that transcript. In the default model of drag our calculation follows recent work from the Levine group exploring the resistance of the elongation complex to rotation^13^, so that the relaxation rate is (in radians/s):

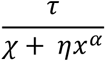

where τ is the net torque experienced by the transcribing RNAP in pN·nm, χ is the DNA twist mobility, set to 0.05 pN·nm·s, η is the rotation viscous drag coefficient, set to 0.0005 pN·nm^1/2^·s, x is the length of the nascent RNA (in nm) attached to the RNAP, and the scaling coefficient α is set to 3/2. In order to avoid unrealistically fast rotational relaxation due to the neglect of viscous drag contributions due to the rotating RNAP itself, we depart from the previous model in replacing the *ηx*^α^ term above with a viscous drag contribution from the polymerase of 0.12 pN·nm·s when the radius of the sphere formed by nascent RNA is less than the RNAP radius (see Supplementary Materials). In addition to using alternative scaling exponents, users can select from alternative models of rotational resistance detailed in the Supplementary Material and documentation included with the *spotter* package.

#### Inclusion of additional topological insulators

Experimental data reported by the Cozzarelli group^59^ indicated that the *E. coli* chromosome is dynamically organized into topological domains that contain roughly 10 kilobase-pairs. The presence of such domains can be mimicked by the inclusion of user-specified barriers located at freely variable positions up- and downstream of a transcription unit. In the simulations reported here we used two approaches. In the first sets of simulations described in Results, topological barriers were omitted entirely, which corresponds to simulating transcription of a linearized template. In the later sets of simulations, topological barriers were placed at −5000 and +5000 of the gene termini to generate a topological domain with an extent approximately equal to the average value identified by the Cozzarelli group.

#### Topoisomerase I and gyrase activity

In addition to responding to RNAP translocation and rotational relaxation, DNA supercoiling also responds to the activity of topoisomerases, whose activity is explicitly modeled in *spotter* simulations. In *E. coli*, the principal topoisomerases are topoisomerase I and gyrase, enzymes that relax negatively- and positively-supercoiled DNA, respectively^34,48^. Briefly, during simulations each enzyme attempts to bind the DNA template at a user-specified rate; the success of binding depends on the steric availability of the randomly-chosen site (determined by the current positions of all RNAPs, the current positions of all other DNA-relaxing enzymes, and the footprint of the enzyme attempting to bind). On successful binding, both types of enzyme alternate between “lag” phases in which they remain bound but in a dormant state and “burst” phases in which they carry out a processive series of relaxation reactions. As is fully detailed in the Supplementary Material, mean dwell times, lag times, burst sizes, and DNA relaxation rates during processive bursts are derived from single-molecule experimental data^34–35,48–51^. On either enzyme’s completing a DNA-relaxation reaction during a *spotter* simulation, supercoiling density is instantaneously equilibrated within the domain where the reaction occurred; the elemental pause entry rate (k_p_) and the rotational relaxation rate for any RNAP acting as a domain boundary for the region in which the enzyme-mediated relaxation occurred are immediately updated to reflect the change in σ. While the parameters describing both topoisomerase I and gyrase activity are both freely variable by the user, the effects of topoisomerases that operate via fundamentally different mechanisms would need to be added in future iterations of the code.

### Translation module

#### Overview

Our goal in developing the *spotter* translation module has been to represent the production of polypeptides from mRNA transcripts at a level of detail sufficient to represent co-transcriptional interactions between ribosomes and RNAPs and to enable meaningful comparison of simulation results with experimental ribosomal profiling data. As has been previously observed^23^, information about the rates of the many subprocesses contributing to ribosomal translocation is not yet clear enough to include in simulation models; in addition, many of the “off-pathway” processes that ribosomes undergo after entering a stalled state remain obscure^24^. Here, we have developed a codon-by-codon model of translational elongation that is resolved into two steps: a consolidated ribosome translocation step (in which the ribosome advances three nucleotides along the mRNA transcript and during which it is assumed that all requisite sub-movements have been accomplished) and a consolidated amino acid addition (in which an amino acid from an cognate tRNA is added to the growing polypeptide and during which it is assumed that non-cognate tRNAs have been rejected). Critically, like the transcription module, and following other models of translation centered on ribo-seq data^21–22^, the translation module is designed to reproduce NGS data: rates for the translocation and amino-acid addition steps at each codon are assigned so that their combined contributions at the position match user-supplied ribosomal profiling dwell times.

#### Assigning rates for translation reactions

At each codon, we divide the unchangeable (ribo-seq derived) net dwell time at that position into contributions from translocation and amino acid addition in a manner intended to be consistent with experimental tRNA abundances in *E. coli*^52^ and a model of tRNA competition reported by Fluitt et al.^53^. Full details of the application of the tRNA competition model appear in the Supplementary Material. Briefly, we first determined the relative usage of each codon using ribo-seq reads from Li et al. for *E. coli* cells growing in rich defined medium^54^ (GEO accession GSE53767), using gene coordinates provided by RegulonDB^42^ in order the match reads to codon identities. Then, using tRNA abundances for *E. coli* under comparable growth conditions^52^, and following the method reported by Fluitt et al., we determined the average arrival time for each tRNA species, from which the average time spent adding an amino acid from the correct cognate tRNA can be calculated (see Supplementary Materials); times required for the addition range from ∼10 to ∼300 ms (Figure 4).

**Figure 4.**
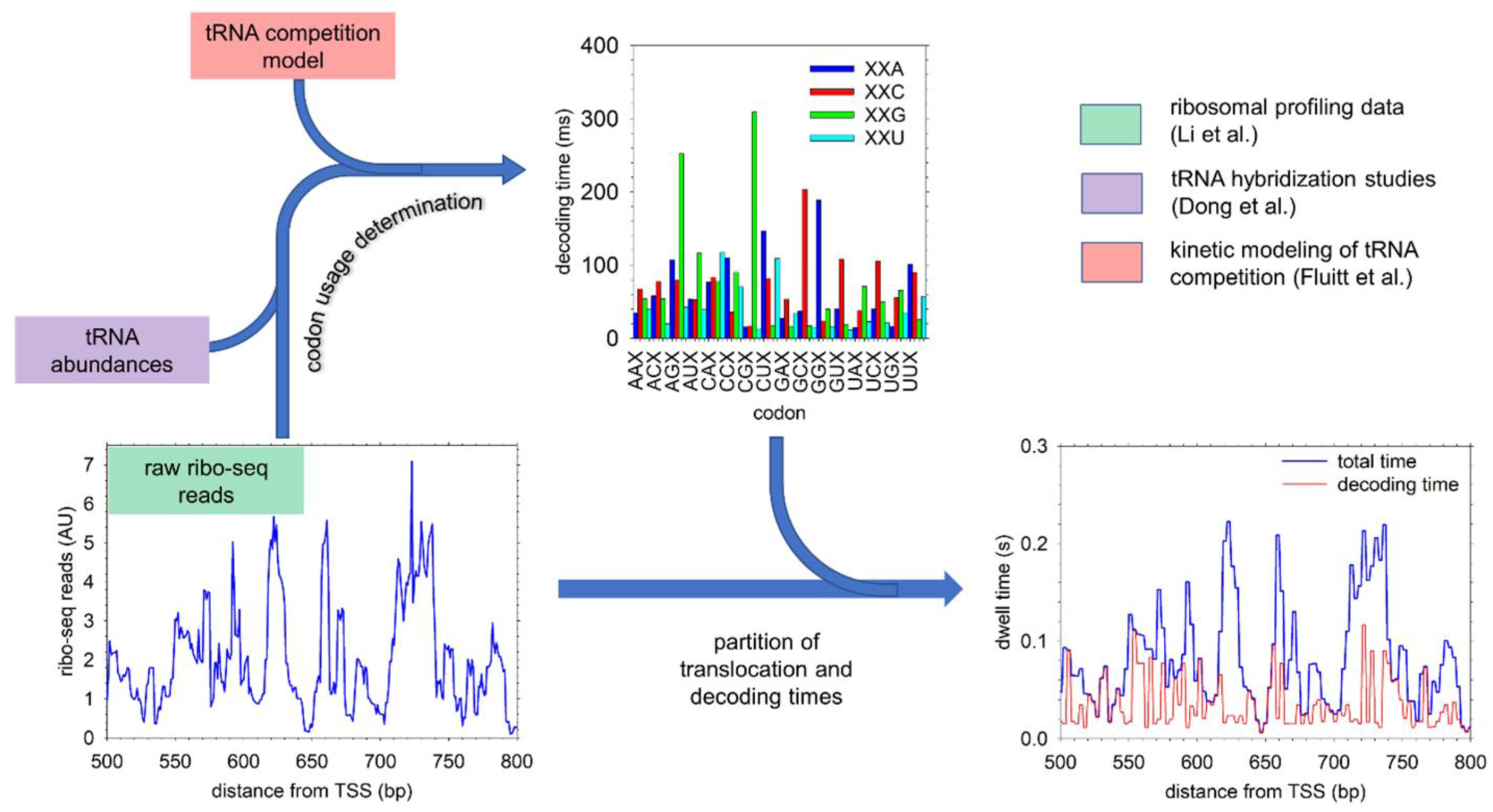
Outline of the translation module. The schematic illustrates experimental information incorporated in the translation module. Decoding times are derived from ribosomal profiling read counts (Li et al.; lower right) for the entire *E. coli* genome, supplemented with tRNA abundance data (Dong et al.). Once whole transcriptome usages were determined for all codons, we used the tRNA competition model described by Fluitt et al. to scale decoding times to match a target global mean translational rate as is detailed in Methods. The scaled decoding times for each codon were then used to partition dwell times obtained from the ribosomal profiling data at every position in the genome. Codon-by-codon total dwell times (blue) and decoding times (red) are plotted at right for a portion of the *gapA* mRNA.

Finally, to partition the fixed (ribo-seq derived) total dwell time into translocation and amino acid addition rates at each position, we assume that competition among tRNAs is relatively invariant at all positions, while translocation rates may be more variable locally. Thus, if the total dwell time at a codon is greater than the average addition time predicted by the tRNA competition model, the amino acid addition time is set to the predicted addition time for that codon and the translocation time is calculated as the total dwell time less the assigned addition time. If the fixed total dwell time is less than the predicted addition time, the translocation time is set to a user-defined minimum (0.1 ms by default), and the amino acid addition time is set to the total dwell time less this minimum. Overall, the model attempts to combine approaches in which dwell times are derived from ribo-seq data^21–22^ with models in which dwell times are derived exclusively from a tRNA competition model^17–19,23^. The requirement that codon-specific dwell times match input sequencing data sometimes precludes the assignment of the full time predicted by the competition model, although in most cases there is sufficient “slack” in the total dwell time to assign an appropriately-low rate for the amino-acid reaction (Figure 4). We also note that, although the examples reported in this paper rely on data specific to *E. coli*, the tRNA competition model we have used is sufficiently flexible to have been used in eukaryotes^17^ and that users interested in creating arbitrary transcriptional landscapes (e.g., with custom-defined pause sites) can easily generate input files appropriate to these systems following instructions included with *spotter* source code.

### Simulation details

Specific details of the different systems are described below, but the following parameters were common to all of the simulations described here: (1) the rotational relaxation rate of the RNAPs was set according to the default model described above (“RNAP rotational relaxation”), in which rotational resistance is related to nascent RNA length with a scaling exponent of 3/2; (2) position-independent rates of elemental pause escape, P2 pause entry and escape, and P3 pause entry and escape were fixed at the values indicated above (“Assignment of rates to RNAP pause-entry and pause-exit reactions”); (3) supercoiling density (σ) within each topological was partitioned into twist and writhe according to in conformity with simulations of DNA minicircles^55^ as described in the Supplementary Material; (4) transcriptional relaxation rates were set to reproduce dwell times calculated from the energy function derived from NET-seq data from Larson et al.^41^. Mean elongation rates over the templates were varied in individual simulations as indicated below.

#### Simulations of lone RNAPs transcribing the rpoB gene

To illustrate *spotter*’s model of transcription, we conducted simulations of isolated (lone) RNAPs transcribing the *E. coli* gene *rpoB* on a template with freely-rotating DNA ends (representing a linear transcriptional template). Hence, although rotation relaxations were included in simulations, in the absence of any domain-defining barriers, the lone RNAPs experienced no net torque during simulations. Relative dwell times were scaled to match the experimental whole-gene mean elongation rate (6.2 nt/s)^39^; differences in dwell times at individual positions spanned approximately two orders of magnitude. To validate our strategy for incorporating experimental NET-seq data, we determined the distribution of dwell times in 1000 simulations; in each simulation RNAPs were allowed to reach a maximum backtracking depth of 0, 10, or 100 bp, and, to match experiment, RNAP positions were recorded at 25 Hz in 4-bp windows. A second set of simulations modeling transcription within an insulated topological domain was conducted with identical parameters, except that the *rpoB* was enclosed within a template with topological insulators located 5000 bp up- and downstream of the transcription start and stop sites, respectively. Topoisomerase I and gyrase were allowed to bind in regions upstream and downstream, respectively, of the gene body at the rates indicated.

#### Simulations of multiple RNAPs transcribing the rpoB gene

To examine the effect of additional traffic on RNAP elongation rates we performed simulations with additional initiation events. Simulations of RNAP pairs designed to test the ability of a trailing RNAP to “rescue” a downstream RNAP trapped in a backtracked pause were identical to those described above for lone RNAPs on linearized templates and on templates enclosed within topological domains, except that on the lead RNAP’s entering a paused state scheduled to last at least 50 s, a second RNAP initiation immediately initiated transcription. Additional simulations included a “release torque threshold” (see “Effects on transcription reactions” in the description of the supercoiling module above) allowing escape from backtracked pause states P2 and P3. Simulations featuring repeated transcriptional initiations are identical to simulations of RNAP pairs within a topological domain using a release torque threshold of 5 pN·nm, except that in these systems transcription is initiated at a rate of 0.1 s^-1^ throughout the simulation and topoisomerase I and gyrase bind at the indicated rates.

#### Simulations of coupled transcription and translation in polycistronic operons

Simulations modeling combined transcription and translation feature the *E. coli gapA*, *dusB-fis*, and *marRAB* operons. With the exception of transcriptional reaction rates specific to their respective sequences, all simulations including translation used parameters identical to those used in simulations of *rpoB* in a closed topological domain with multiple transcriptional initiation events and a release torque threshold of 5 pN·nm; the transcriptional initiation rate was set 0.1 s^-1^ in all simulations of coupled transcription and translation. Translation initiation rates for the *gapA* operon were set as indicated; translation initiation rates for *dusB-fis* and *marRAB* were set based on translation efficiencies reported by Li et al.^54^.

### Analysis and visualization of simulations

*spotter* automatically accumulates the output from many replicate simulation trajectories to give summaries of RNA and protein production statistics, thereby providing a route to making direct comparisons with cellular-scale experimental data. But *spotter* is also intended to give detailed output data characterizing individual simulation trajectories. To this end, *spotter* generates single-molecule traces, dwell-time distributions, simulated sequencing data, and kymographs that illustrate changes in local supercoiling density over the course of a trajectory. In addition, and perhaps most useful, *spotter* is equipped with a visualization toolkit that automatically generates two different molecular-level views of the simulated behavior.

## RESULTS

### Simulations of transcription by lone RNAPs reproduce single-molecule and NGS data

To provide a first illustration of *spotter*’s capabilities, we focus on the modeling of transcription alone. We start with straightforward simulations of lone RNAPs on the *E. coli* gene *rpoB*, a gene that we chose as it was recently the subject of single-molecule experiments that provided a detailed characterization of RNAP pausing and release.^39^ Since these *in vitro* experiments did not include any translational machinery, all translational reactions were omitted. As primary input to the simulations, we use gene-position-specific dwell times that we derive from the NET-seq data for the *rpoB* gene reported by Larson et al. (see Methods). In all of the simulations shown here, the RNAP is allowed to enter paused states and backtracking is enabled, but separate sets of simulations were performed in which the maximum allowed depth of backtracking was varied from 100 bp to 10 bp to 0 bp (i.e. no backtracking). In a representative trace from one of these simulations (Figure 5A), the mean elongation rate over a 200 s interval can be seen to match the expected whole-gene elongation rate of 6.2 nt/s derived from experiment^39^. On a shorter timescale (5-20 s), however, the elongation rate can be seen to vary widely, and at least two extended backtracking events can be observed: in the enlarged view of these events shown at right, we see that the position of the RNAP (blue line) moves backwards to upstream positions in the gene, while the 3’ end of the RNA 3’ (red line) remains unchanged. Both backtracking events are ultimately ended when the RNA is subjected to a chance processing event that cleaves the RNA: this can be seen to occur when the length of the RNA (indicated by the 3’ position) abruptly drops (vertical red lines). Importantly, aggregating the data from 1000 simulations confirms the correct implementation of *spotter*: the distribution of dwell times experienced by the RNAP is in excellent agreement with the corresponding distribution obtained from single-molecule experiments performed by the Dekker group (Figure 5B), and the simulated NET-seq data are in similarly good agreement with the experimentally-derived RNAP occupancy profile across the *rpoB* gene reported by the Greenleaf, Landick, and Weissman groups (Figure 5C). Interestingly, the level of agreement between simulation and both types of experiment appears to be insensitive to the maximum backtracking depth permitted in the simulations (Figure 5C). Overall then, these results indicate that *spotter* can be effectively parameterized so that it produces behavior that is simultaneously consistent with single-molecule data and with whole-cell NGS data such as NET-seq.

**Figure 5.**
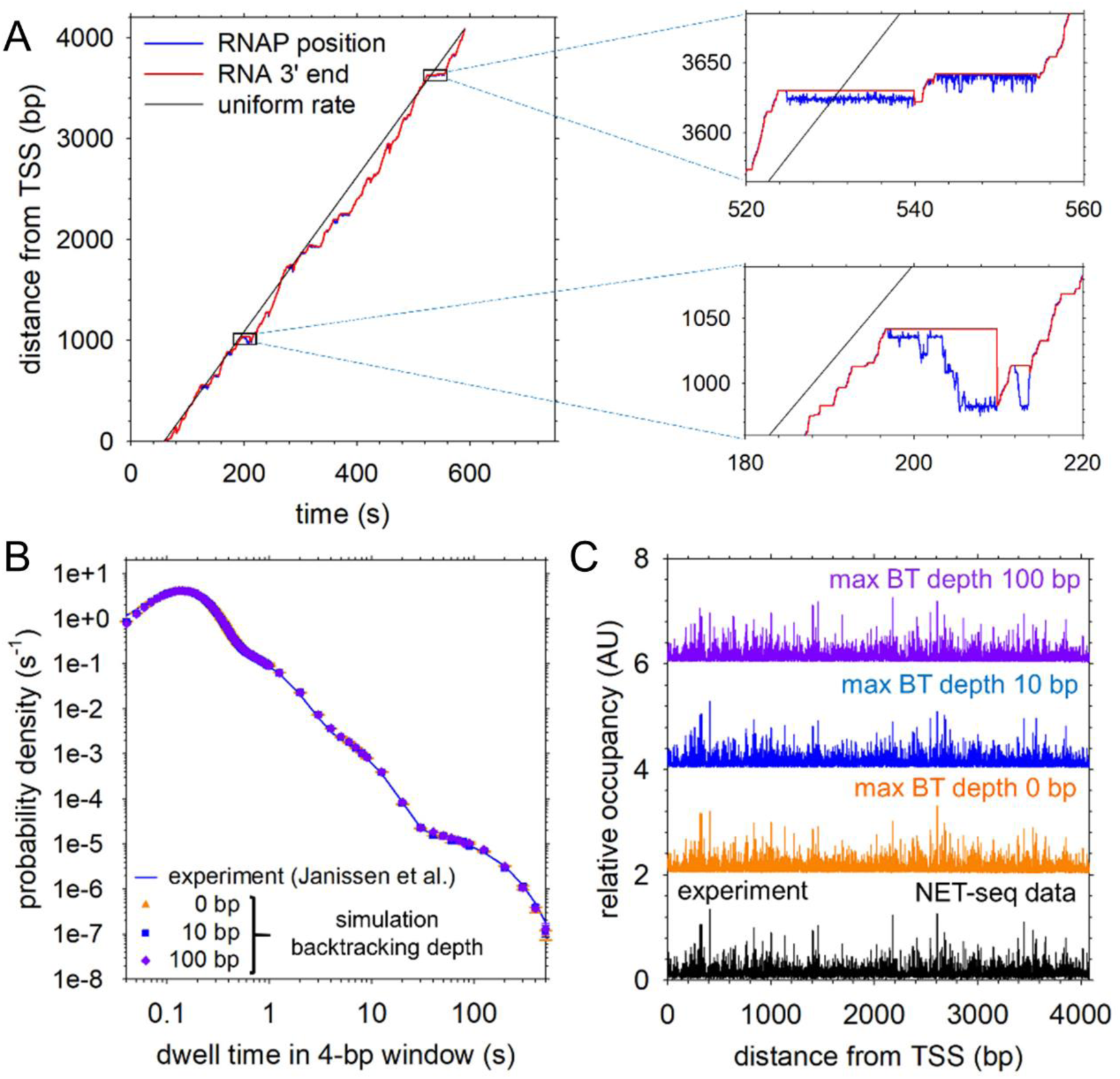
Validation of the transcription module. **A.** Representative single-molecule trace of RNAP progress along the *rpoB* gene in simulations in which RNAPs were allowed a maximum backtracking depth of 100 bp from the position in which they entered an advanced pause (state P2 or P3). The model was set to match an experimentally-observed mean elongation rate (6.2 nt/s) from single-molecule studies of transcription on *rpoB*^30^. For comparison, dashed lines represent the progress of an idealized RNAP transcribing *rpoB* at the same rate as the simulated RNAP but advancing through all template positions at an identical rate. Insets depict 40-second observation windows from the trace. **B**. Comparison of experimental dwell-time distribution and the dwell-time distributions in simulations using the *spotter* transcriptional model with varying backtracking depths. The solid blue line is the fit to experimental data^30^ for a 4-bp observation window as detailed in Methods. **C.** Comparison of simulated and NET-seq derived RNAP occupancy. Positions of nascent RNA 3’ ends in 1000 were determined at 0.04 s intervals. Each simulation consisted of a single transcriptional run for an isolated RNAP. RNAPs were permitted the maximum backtracking depths listed. Peak height at each position represents the relative number of 3’ reads for that position in the 1000-simulation aggregate. The solid black line represents relative occupancy derived from NET-seq reads across the *E. coli* genome as described by Larson et al. and described in Methods.

### A supercoiling-mediated traffic jam of two RNAPs

In the above simulations we considered only the progress of isolated RNAPs, and backtracked RNAPs were able to resume active transcription only upon cleavage of their extruded 3’ RNA; this model is consistent with the Dekker group’s single-molecule experiments which focused on lone RNAPs.^39^ But transcription units *in vivo* are very frequently occupied by multiple RNAPs producing copies of the same transcript, and a growing body of evidence suggests that communication between these RNAPs is mediated by supercoiling of the intervening DNA.^16,47.56^ Here, we provide a simple illustration of *spotter*’s ability to model this interplay between transcription and supercoiling, by modeling the combined transcription of two RNAPs on the *rpoB* gene. Ultimately, we use the model to propose a mechanism to explain the puzzling observation that while RNAP backtracking appears to be ubiquitous in lone-RNAP experiments performed *in vitro*,^39–40^ it appears to be relatively rare in highly-transcribed operons *in vivo*, at least as suggested by cellular-scale NET-seq data.^41^

To isolate an interesting scenario, we returned to the above simulations of lone RNAPs transcribing the *rpoB* template, and selected for further study a *spotter* trajectory in which the RNAP was observed to enter a long-lived backtracked pause state. A kymograph illustrating the progress of the RNAP in this specific trajectory is shown in Figure 6A: the RNAP begins transcription at t = 9 s, transcribes successfully for ∼40 s, before entering a paused state at template position 412 that, in turn, leads to a backtracked state that persists until a chance RNA cleavage event returns it to on-pathway transcription.

**Figure 6.**
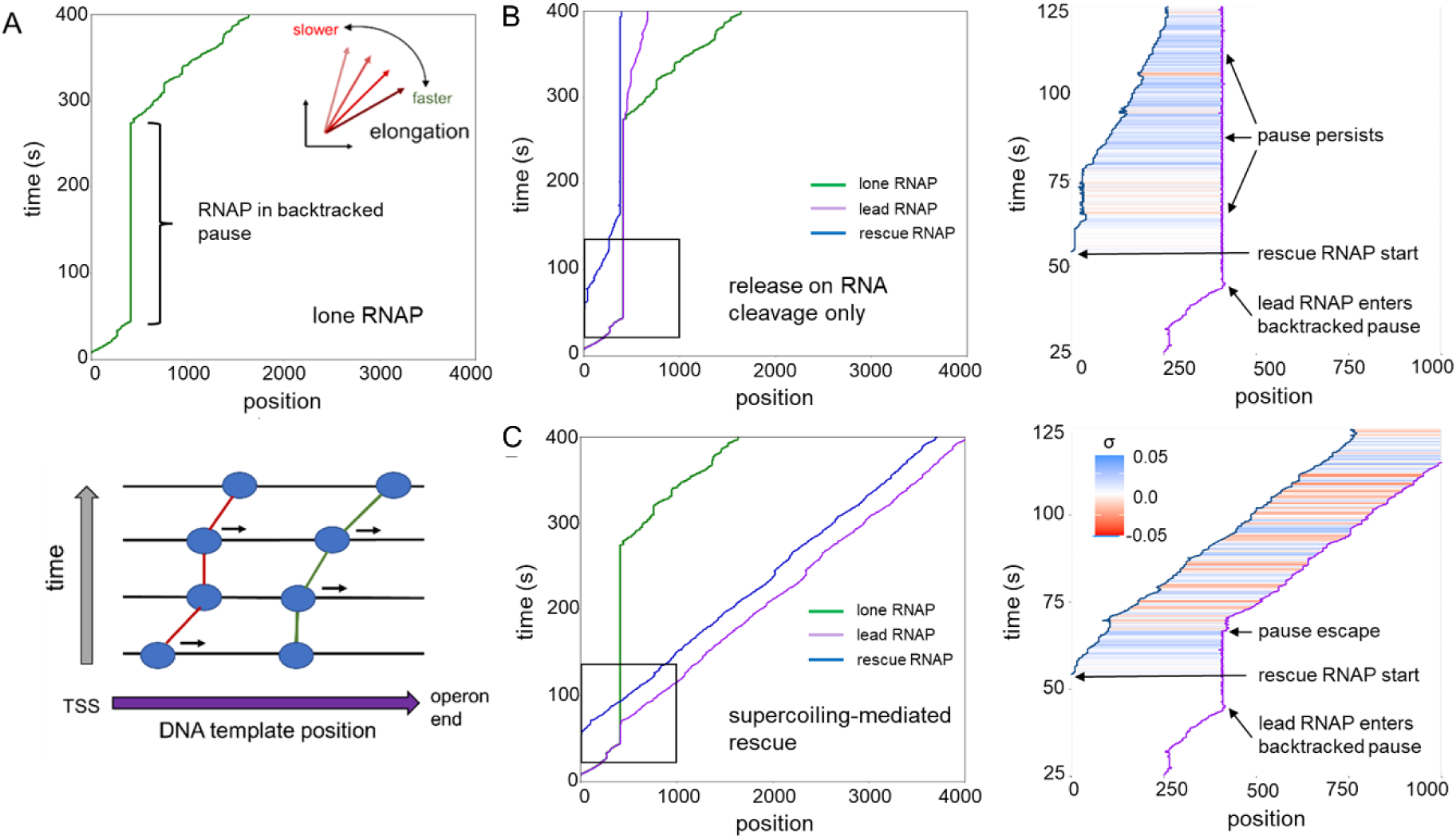
Supercoiling-mediated rescue of a backtracked RNAP. **A**. Kymograph representing the progress of an isolated RNAP. Progress along the DNA template is represented by rightward movement along the x-axis (bottom panel); stalled RNAPs appear as vertical lines. The trace was produced from a simulation of a single RNAP transcribing *rpoB* from a template with freely-rotating DNA ends and a relative rotational relaxation rate set as described in Methods; all other reaction rates were set to the values used in Figure 4. For clarity, only the position of the mRNA 3’ end is plotted. **B.** Rescue of a stalled RNAP. The simulation for which the traces are plotted was identical to the lone-RNAP simulation at left except that a second “rescue” RNAP initiated transcription 0.001 s after the lead RNAP entered the long-duration pause at position 412. When subject to 5 pN·nm of torque, an RNAP in advanced pause state P2 or P3 at the site of initial pause is released into the pretranslocated state. Positions of the lead (purple) and rescue (blue) RNAP are plotted; the lone-RNAP trace (green) is replotted for comparison. At right, local supercoiling density (σ) is plotted for the region boxed at left: inter-RNAP σ ranges alternates between positive and negative values as the lead and rescue RNAP “pull” and “push” one another, respectively. σ is plotted at 5 bp ˣ 0.1 s resolution. **C.** Rescue depends on supercoiling-assisted release from advanced pause states. RNAP positions are plotted for a simulation identical to that represented in B., except that RNAPs in state P2 or P3 can only return to the pretranslocated state through cleavage of extruded 3’ RNA; rates of back- and forward-tracking in these states are affected by supercoiling, but pause duration is not.

To ask how this behavior might change in the presence of additional traffic, we repeated exactly the simulation of this trajectory but with the introduction of a second RNAP that was programmed to initiate transcription immediately after the leading RNAP entered its paused state. A kymograph illustrating what happens is shown in Figure 6B. The second (lagging) RNAP falls prey to a transient pause at template position 11, and when it emerges from this pause its elongation rate is clearly reduced from the rate displayed by the lone RNAPs shown in Figure 5. This effect can be traced to the supercoiling that builds up in the inter-RNAP region (positive σ): the inability of the leading RNAP to exit its paused state means that every forward move of the lagging RNAP is accompanied by an increase in positive supercoiling between the two RNAPs, and further progress is only possible because of slow rotational relaxation of both RNAPs. Eventually, the lagging RNAP “catches up” with the leading RNAP and both become mired in paused states. The chance escape of the leading RNAP from its paused state allows it to continue translocation, but since the lagging RNAP is now paused, we see the same slowing of the elongation rate that was previously seen for the lagging RNAP, only now the supercoiling in the inter-RNAP region is predominantly negative. Still later, the lagging RNAP exits its paused state thanks to the chance cleavage of the extruded RNA, and from this point on both RNAPs cooperate in the push-pull fashion seen above, with elongation finally completing after ∼1200 s.

### A torque-mediated pause-release mechanism relieves a traffic jam

The above results suggest that the combination of supercoiling-mediated RNAP-RNAP communication and long-lived paused states could, in principle, lead to catastrophic transcriptional traffic jams *in vivo*. One proposed mechanism for the avoidance of such traffic jams involves transcriptional “backstopping,” in which a trailing RNAP on the DNA template or a co-transcriptionally translating ribosome on the mRNA template encourages transcriptional progress by sterically preventing RNAP backtracking^57–58^. While the presence of such transcriptional backstops can effectively restrict the movement of an RNAP trapped in a long-lived pause state (allowing the RNAP to regain the position at which it first entered the pause more rapidly), single molecule data suggest that return to the site of pause entry is insufficient to promote the transition to on-pathway transcription^39–40,46^. As another potential contributor to the relief of transcriptional traffic jams, we propose here the possibility that a trailing RNAP might release (“rescue”) a leading paused RNAP by giving it a DNA-supercoiling-mediated “push” once it has returned (by stochastic translocation) to the site of its initial pause. To model this possibility, we allow RNAPs in extended pause states to be driven to return to their pretranslocated state if the supercoiling-mediated torque that they experience exceeds a threshold that we term the “release torque threshold” (see Methods above). Viewed from this perspective, the above simulations, in which such a mechanism was disallowed, correspond to ones in which the release torque threshold was set to infinity.

To explore the possible consequences of introducing such a mechanism, we repeated exactly the simulation of this trajectory, but with the hypothetical release torque threshold now set to a more realizable value of 5 pN·nm. A kymograph illustrating what happens is shown in Figure 6C: the lagging RNAP transcribes successfully to template position 99, there is a brief period (∼1s) during which both RNAPs appear to suspend transcription, before the leading RNAP is awoken from its paused state, and both RNAPs resume elongation. The duration of the leading RNAP’s backtracked pause is therefore reduced from ∼200 seconds in the lone-RNAP simulation to less than 50 seconds, and both lead and rescue RNAPs are able to complete transcription of the *rpoB* template is less than 400 seconds (Figure 6C, left). Strikingly, once the leading RNAP resumes productive transcription, the two RNAPs maintain a relatively stable separation of ∼300 bp. The source of this effect is apparent in the accompanying plot of supercoiling density (Figure 6C, right), where negative inter-RNAP σ alternates with positive inter-RNAP σ as the leading or the lagging RNAP, respectively, outstrips its counterpart. This pattern is consistent with predictions made by the Gedeon and Jacobs-Wagner groups of supercoiling-mediated “pulling” of upstream RNAPs and “pushing” of downstream RNAPs by an actively-transcribing RNAP^9,16^.

### Transcription of a lone RNAP within an extended topological domain is sensitive to topoisomerases

For simplicity, the simulations above model transcription on a DNA template whose ends are assumed to be free to rotate, resulting in supercoiling densities of zero in the regions of the gene downstream of the leading RNAP and upstream of the lagging RNAP. While linearized templates like this can be important for comparison with *in vitro* experiments, transcription *in vivo* (at least in *E. coli*) is thought to occur within topologically-insulated domains whose mean length is ∼10 kilobase-pairs^59^. Because transcription within a closed domain results in the accumulation of negative supercoiling upstream of the transcription start site (TSS) and positive supercoiling downstream of the end of the template, it requires the activity of enzymes that can relax template DNA in these regions. In order to highlight specifically *spotter*’s ability to model the effects of these topoisomerases, we first return to simulations of lone RNAPs transcribing the *rpoB* template but we now situate it at the center of a 10 kilobase-pair topological domain. To allow the contributions of DNA-relaxing processes to transcriptional dynamics to be observed we allowed topoisomerase I to bind upstream of the ∼4000-bp *rpoB* gene and allowed gyrase to bind downstream of the gene (Figure 7A, top); in this scheme, each enzyme binds in the region where it can effectively support translation, with topoisomerase I relaxing the negative supercoils that accumulate upstream and gyrase relaxing the positive supercoiling relaxing the positive supercoils that accumulate downstream of RNAPs advancing within an insulated domain.

**Figure 7.**
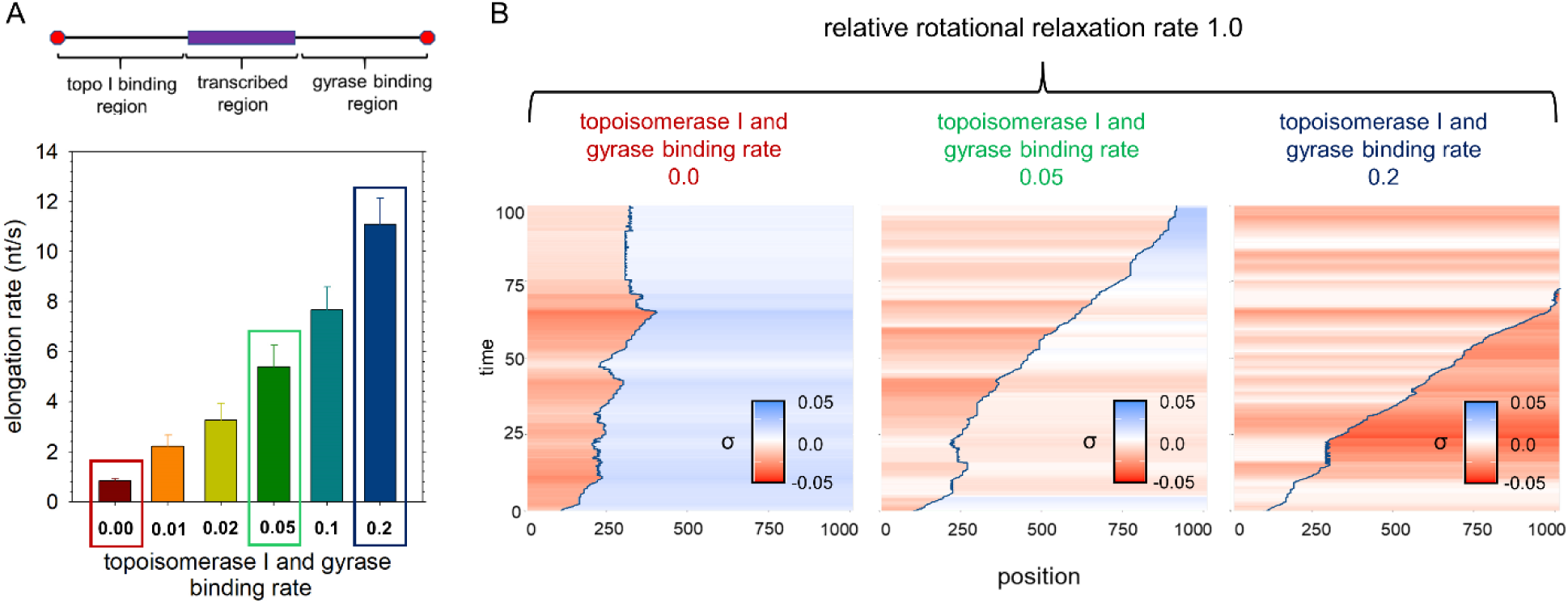
Transcriptional elongation rates are highly sensitive to processes relaxing supercoiled DNA. **A.** Elongation rates are affected by rates of enzymatic and RNAP-rotation-driven relaxation of DNA. Mean elongation rates from simulations of lone RNAPs transcribing *rpoB* within a topologically-insulated 10 kbp domain are plotted. Rates of topoisomerase I and gyrase binding were varied as indicated; results from 1000 simulations for each condition are shown (error bars indicate standard deviations). Listed topoisomerase I and gyrase binding rates are the rates (per second) at which each enzyme attempts to bind the 10 kbp DNA template. To facilitate transcription in the simulations, topoisomerase I was allowed to bind only the region upstream of the TSS, where it works to relax negatively-supercoiled DNA, and gyrase was allowed to bind only to the region downstream of the *rpoB* gene, where it works to relax positively-supercoiled template DNA (schematic at top). **B.** Representative kymographs illustrating enzyme effects on supercoiling density. Plots under the red, green, and blue headings correspond to the boxed regions in A. At low rates of enzymatic activity (left) RNAP progress is slow, with accumulation of negative supercoiling downstream of the transcribing RNAP. As rates of enzymatic activity increase (middle), the accumulation of downstream negative supercoiling is reduced, with elongation rates rising accordingly. At high levels of enzymatic activity (right), σ downstream of the transcribing RNAP is negative, reflecting the ability of gyrase to introduce negative supercoiling, in contrast with topoisomerase I, which relaxes negative supercoiling but cannot produce positive supercoiling.

The mean elongation rates of lone RNAPs transcribing within the topological domain are found to be strongly influenced by topoisomerases, dropping from a maximum of 11 nt/s to 1 nt/s as the binding rates of the two enzymes change from 0.2 to 0 attempts to bind their respective template regions per second. While it is to be expected that these effects might change somewhat depending on how fast one assumes that RNAP rotational relaxation occurs, it is to be noted that at sufficiently high topoisomerase activity, the mean elongation rate (11 nt/s) is predicted to exceed that observed on a linearized (i.e. topologically unconstrained) template (6.2 nt/s; see earlier). The origins of this observation can be traced to Figure 7B which shows how σ varies across the *rpoB* part of the topological domain. When the binding rates of topoisomerase I and gyrase are highest (right-most panel of Figure 7B), the downstream σ (to the right of the RNAP position) is consistently more negative than the upstream σ (to the left of the RNAP position). This reflects the known differences in the behavior of gyrase, which binds in the downstream region and is capable of changing σ from positive to negative, and topoisomerase I, which binds in the upstream region and is capable only of returning σ from negative values to zero. The origins of the slowdown in elongation rate that occurs in the absence of topoisomerases is similarly apparent from the left-most panel of Figure 7B. In this case, the upstream and downstream σ values remain stubbornly positive and negative, respectively, and repeated pausing and backtracking events are observed.

### Topoisomerases allow efficient transcription to be maintained even at high transcription initiation rates

As a final illustration of the *spotter* supercoiling model, we modeled the effects of additional transcriptional traffic in simulations occurring within the same topological domain. First, we assessed again the ability of a second, lagging RNAP to rescue a leading RNAP that had entered a backtracked pause during transcription within the topological domain. As is illustrated in Figures 8A and B, a second RNAP initiating immediately after the moment that the lead RNAP enters a long-duration pause can release the lead RNAP from the backtracked state, but the difference in mean elongation rates between lone and rescued RNAPs is more modest than was seen in simulations of transcription on a template with freely-rotating ends (see earlier).

**Figure 8.**
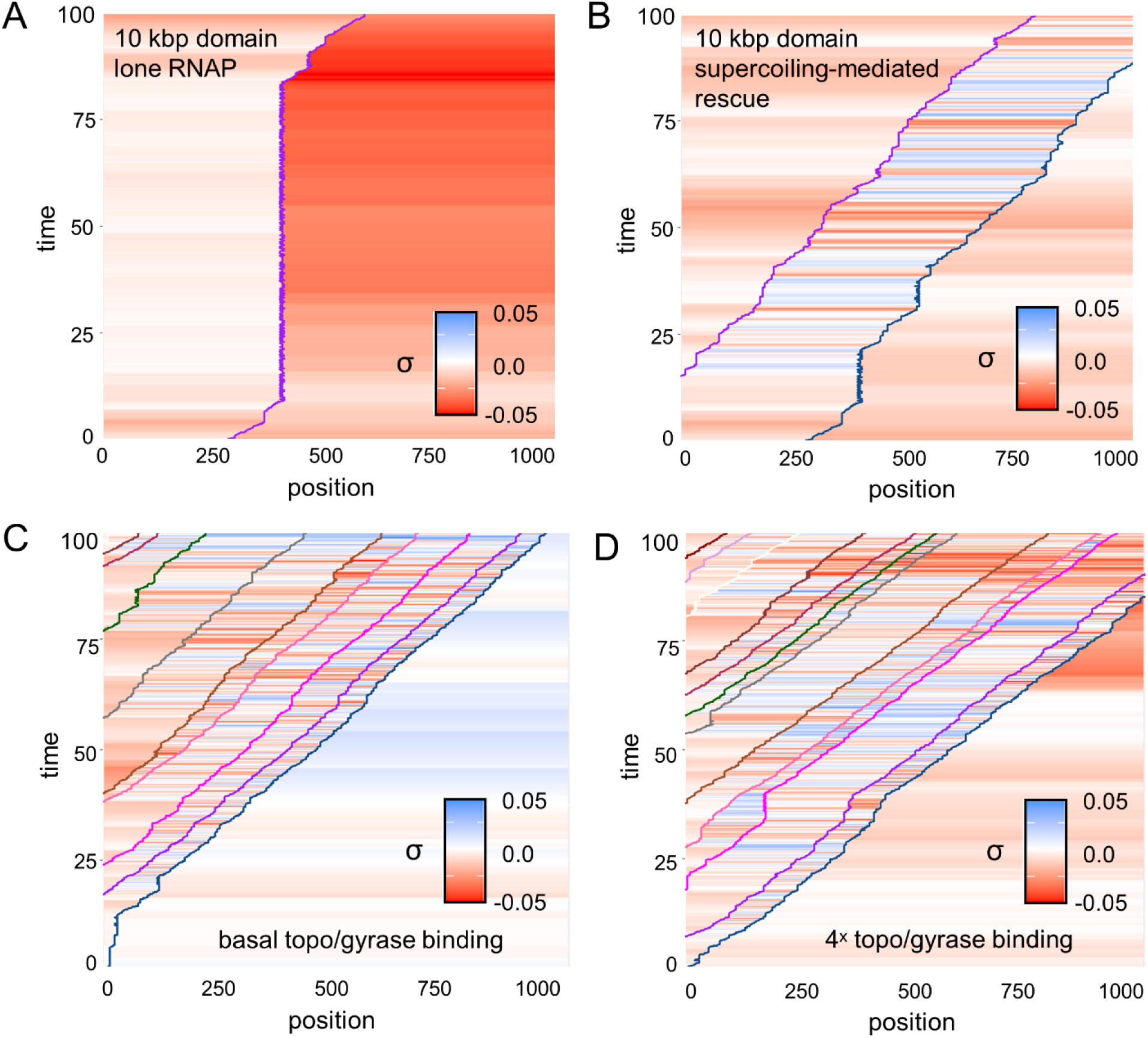
Rescue of a stalled RNAP within a topological domain. **A**. Kymograph illustrating an RNAP experiencing a long-duration pause with backtracking while transcribing *rpoB*. The transcribed sequence and insulated topological domain used in the simulation are identical to those described in Figure 7. The relative RNAP rotational relaxation rate was set to 1.0 and rates of binding for both topoisomerase I and gyrase were set to 0.0001 bp^-1^ s^-1^. Supercoiling density (σ) is depicted at 5 bp ˣ 0.1 second resolution. **B.** Rescue of the stalled RNAP. The simulation for which the kymograph is plotted was run identically to the lone-RNAP simulation in A. except that a second “rescue” RNAP initiated transcription 0.001 s after the lead RNAP entered the long-duration pause at 9 s. **C.** Inter-RNAP supercoiling density in transcriptional traffic. A kymograph representing multiple RNAPs transcribing *rpoB* concurrently is plotted. The transcribed sequence and insulted topological domain used in the simulation are identical to those used in A.; relative RNA rotational relaxation rates were set to 1.0; topoisomerase I and gyrase binding rates were set to 0.5 attempts on their binding regions per second; and the assistive torque required to release an RNAP in pause state P2 or P3 was set to 5 pN·nm. The rate of RNAP loading attempts was set to a mean of 0.1 s^-1^. **D.** Same as C., but the rates of topoisomerase I and gyrase binding were increased four-fold to 0.2 attempts on each binding region per second.

Next, we simulated transcription in much heavier traffic, i.e. with many more RNAPs present. As a demonstration of *spotter*’s ability to monitor local supercoiling density at high but realistic RNAP initiation frequencies, we performed simulations in which RNAPs initiated at a rate of 0.1 s^-1^. With basal-level topoisomerase I and gyrase binding rates (Figure 8C), the mean elongation rate is 10.4 ± 1.4 nt/s, and the RNAPs again move in the push-pull mode, maintaining roughly their initial separation distances along the template. With four-fold higher binding rates, the mean elongation rate rises till further to 12.1 ± 1.6 nt/s. Although the results we present here are intended only to illustrate the capabilities of the simulation model, together they suggest that variations in transcription elongation rates depend crucially on the complicated interplay of RNAP-rotation-driven and enzymatic relaxation, rates of RNAP pause entry and escape, and transcriptional initiation rates. The ability of *spotter* to integrate these processes in a single simulation model is thus likely to be a valuable resource in efforts to investigate the role of DNA supercoiling in modulating transcription.

### Illustration of the *spotter* translation module

Finally, to illustrate the *spotter* translational model, we consider simulations in which supercoiling-mediated transcription and translation are both simultaneously modeled. We begin with simulations of the monocistronic *gapA* operon from *E. coli*. For transcription, we again use as input position-specific dwell times that we derive from the NET-seq data for the *gapA* gene reported by Larson et al. For translation, we use as input position-specific dwell times that we derive from ribosomal profiling (ribo-seq) data reported for the same gene by Li et al.^54^; in our model, these dwell times are partitioned into separate contributions from ribosome translocation and amino acid addition steps. Separate sets of simulations were performed in which the mean rate of translation initiation was varied from 0.05 to 0.1 to 0.5 s^-1^. Encouragingly, the simulated and experimental ribo-seq data were in excellent agreement regardless of the assigned translation initiation rate (Figure 9A), and since the latter rate is even higher than the upper range of experimentally measured initiation rates (∼0.3 s^-1(ref^ ^60)^) there appears to be no need to invoke additional mechanisms of inter-ribosomal communication in order to match experiment using physiologically-reasonable polypeptide elongation rates (occurring at a median rate of 12.5 codons/s^61^).

**Figure 9.**
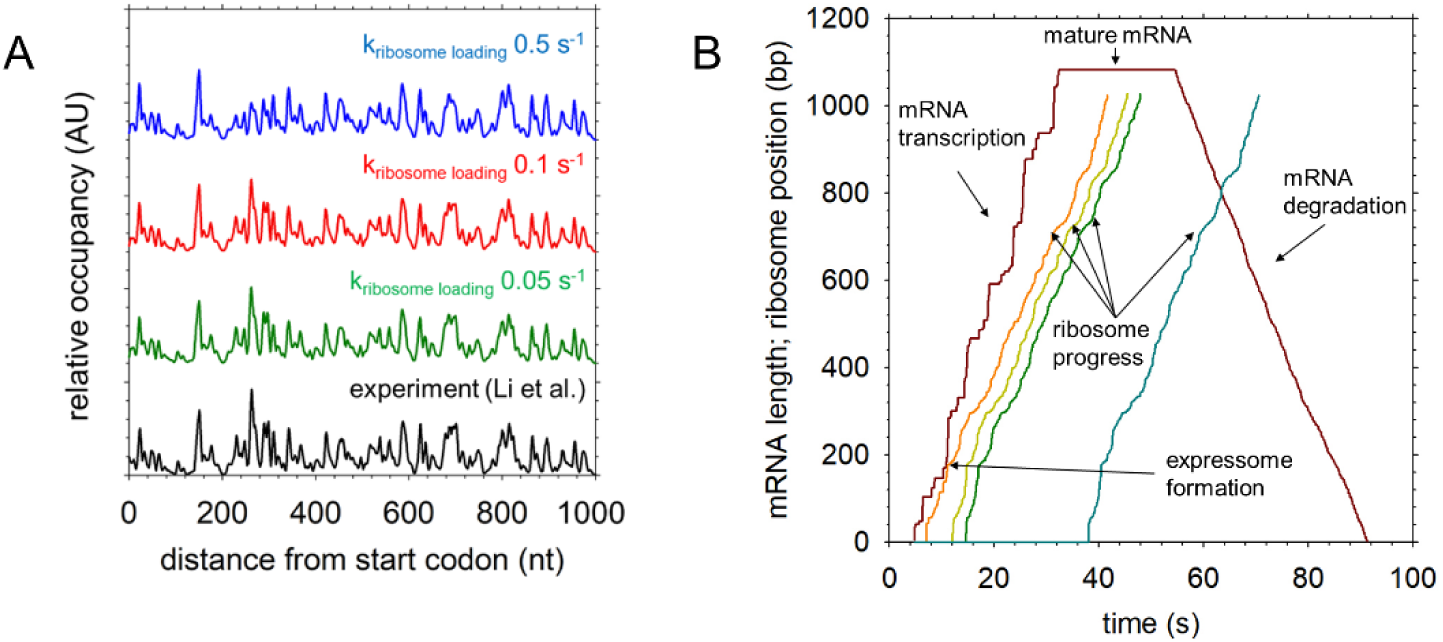
Illustration of the *spotter* translation module. **A**. Validation of *spotter* ribosomal profiling. Ribosome profiling data for *gapA* from Li et al. are plotted in black and compared with simulation ribosomal occupancy in systems employing the one-state model with rates set to produce a mean elongation rate for lone RNAPs of 40 nt/s over the transcription unit on a linearized plasmid template and a transcription initiation rate of 0.1 s^-1^; the mean translation elongation rate over the *gapA* gene was set to 40 nt/s (13.33 codons/s) with translation initiation rates set to the listed values. 1000 15-minute simulations were run for each condition. **B.** *spotter* representation of a complete mRNA lifecycle. By plotting RNA length (dark red line) and ribosomal positions (orange, yellow, green, and blue lines) using the same axis, *spotter* illustrates inter-ribosomal spacing and expressome formation. In the nascent mRNA (the region over which RNA length is increasing), the RNA length plot also represents RNAP position, so that the formation and dissolution of expressomes can easily be visualized; plots can also illustrate simultaneous translation and degradation, which occurs in this plot in the continued progress of the ribosome represented by the blue line after degradation is initiated at ∼60 seconds. The example plotted used parameters identical to those used for the red line in A., except that mean elongation rates for both transcription and translation were set to 30 nt/s (10 codons/s).

The interplay between the transcribing RNAP and the ribosomes that are attached to its nascent mRNA can be visualized using kymographs of the kind shown in Figure 9B. Here, the dark red line plots the transcription of the mRNA, which was modeled in the same way used in the earlier Figures: the line rises stochastically until the point at which transcription is complete and the full-length mRNA is produced; some time later, the mRNA is engaged (again, stochastically) by the degradation machinery, which acts from the 5’ end, and the length of the mRNA begins to decrease. The adjacent three colored lines (orange, yellow, green) show the progress of three ribosomes that begin translation when the mRNA is still nascent, and that are therefore engaged in co-transcriptional translation. The final line (cyan) shows the progress of a ribosome that only begins translation after the mature mRNA has been fully transcribed, i.e. post-transcriptional translation. Two points on the kymograph are especially notable. The first, occurring at ∼12 s, shows the brief formation of an expressome as the point at which the line representing mRNA transcription and the line representing the first (pioneer) ribosome come into contact. In this particular simulation trajectory, the formation of the expressome is a short-lived event, as no energetic reward is assigned to its formation and the mean rate of transcription comfortably exceeds the corresponding rate of translation; this behavior would change if the user desired. The second event of interest is the crossing of the mRNA degradation line and the translation progress of the finally-loaded ribosome. This is indicative of a “foot race” between the final ribosome and the mRNA degradation machinery.

Two important features of *spotter* that we have yet to touch on are: (a) its ability to handle translation from polycistronic operons, and (b) its use in automatically generating scripts that allow transcription and translation to be visualized in molecular terms. To illustrate both of these features we performed simulations of two more *E. coli* operons: the bicistronic *dusB-fis* operon and the tricistronic *marRAB* operons. The *dusB-fis* operon is unusual in that the experimentally measured translation efficiencies of its two genes differ drastically: in rich-medium growth conditions, the *fis* gene is translated ∼150x more than the *dusB* gene. This behavior, which is modeled in *spotter* simulations by assigning the *fis* gene a translation initiation rate that is 150x higher than that assigned to the *dusB* gene, leads to simulation snapshots such as that shown in Figure 10A. Here we see the *dusB-fis* gene at the left, with 8 actively transcribing RNAPs (blue spheres) on it, and with their nascent mRNAs extending upwards; we see that none of the nascent mRNAs are being translated. Adjacent to the gene are mature *dusB-fis* mRNAs, arrayed in the order that they were synthesized, with the oldest mRNAs at the far right of the panel. In each mature mRNA, the *dusB* gene is colored in cyan and the fis gene is red; untranslated regions of the mRNA are colored white, with the 5’-untranslated region (UTR) shown at the top, the 3’-UTR at the bottom, and a short inter-genic region shown in between. The different levels of translational activity on the two parts of the mRNA are immediately obvious.

**Figure 10.**
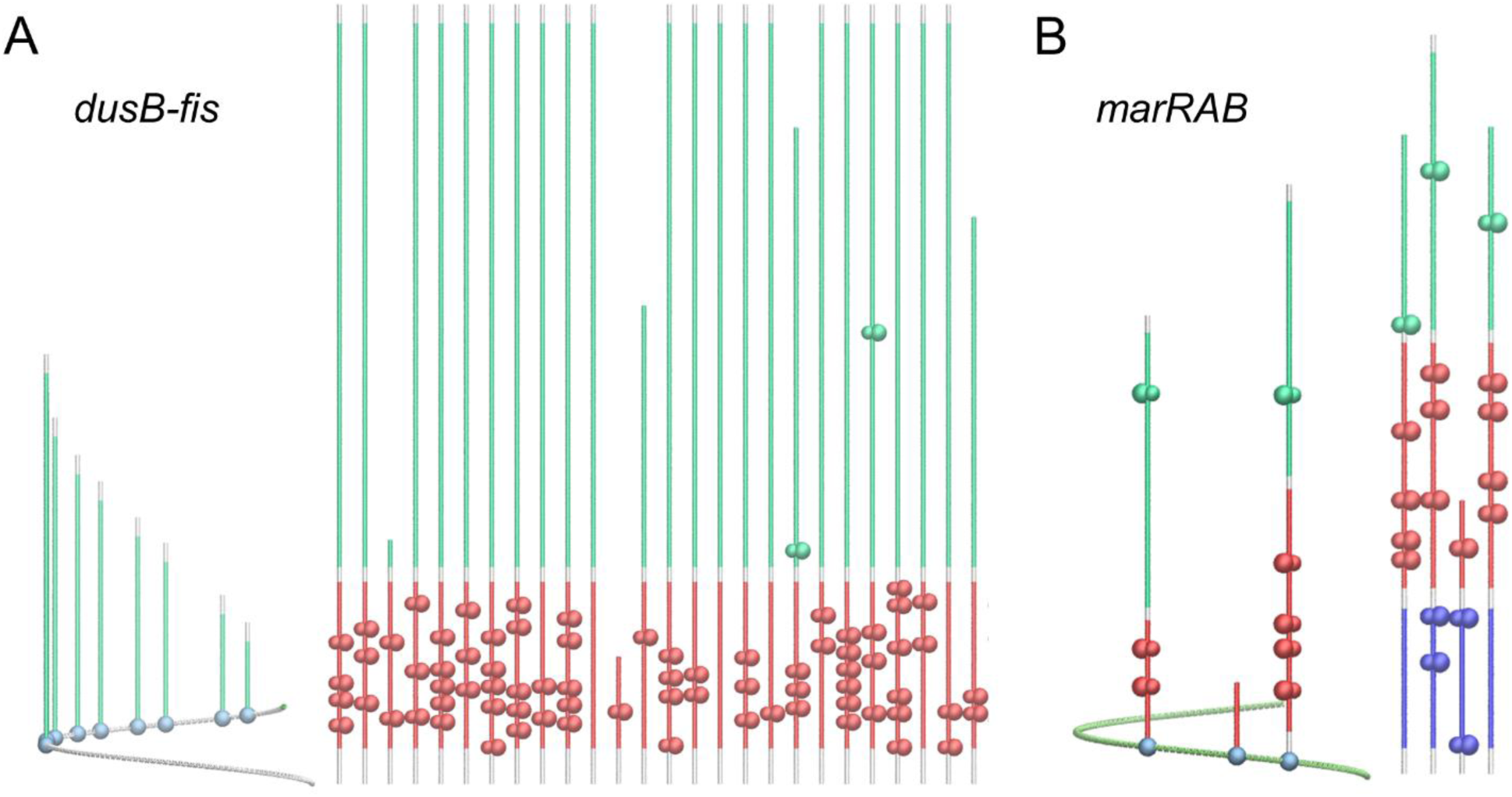
Polycistronic transcription units. **A.** Snapshot from a movie representing transcription and translation on the bicistronic *E. coli* operon, *dusB-fis*, where the translation initiation rate of the downstream gene (*fis*, red gene and ribosomes) is ∼100-fold greater than that of the upstream gene (*dusB*, green gene and ribosomes). Movies for polycistronic operons can be made with a single VMD command using scripts automatically generated during simulations. **B.** Snapshot from a movie representing transcription and translation on the tricistronic *E. coli* operon *marRAB*. Complete movies for A. and B. are included in Supplementary Materials as Movie S4 and Movie S5.

The *marRAB* operon contains genes whose translation efficiencies are more evenly matched: according to the data of Li et al.^54^, the relative rates of translation for *marR*, *marA* and *marB* are 0.04 s^-1^, 0.3 s^-1^ and 0.07 s^-1^, and the snapshot shown in Figure 10B reflects this. We note also that one of the three RNAPs caught in the act of transcribing the operon is undergoing cotranscriptional degradation of its message. Since *spotter* reports residue-level information about the positions of ribosomes and RNAPs throughout an entire simulation trajectory, it provides straightforward opportunities to assess the extent to which phenomena such as the following occur: (a) cotranscriptional translation, (b) expressome formation, (c) cotranscriptional degradation.

## DISCUSSION

*spotter* is designed to extend significantly the range of systems that can be represented in stochastic simulations of transcription and translation in prokaryotes. Challenges arising from the need to manage many interconnected reactions in stochastic simulation algorithms have in the past led highly-sophisticated, sequence-dependent stochastic models to focus on either RNA production or protein production, treating only one of the three modules that are included in *spotter* in isolation. On the other hand, simulation models that have been more ambitious in the number of processes they include have typically relied on simplified representations of these processes, limiting the number of reactions available to RNAPs, ribosomes, and DNA-relaxing enzymes. *spotter*, in contrast, aims to offer highly-detailed models of RNAP transcript elongation, DNA supercoiling, and protein production, providing the simulation machinery necessary to thoroughly investigate the still somewhat obscure relationship between translation and transcriptional progress^58^. Moreover, in its ability to merge models of supercoiling and RNAP backtracking, *spotter* offers opportunities to bring together simulation methods derived from: (a) single-molecule studies characterizing backtracking^3–4,7–8,40^ and (b) single-molecule studies characterizing supercoiling.^9,13–14^

Many components of the *spotter* machinery follow strategies developed by others, and it is important, therefore, to be clear about which of its components are, to our knowledge, novel, lest readers leave with the impression that *spotter* simply cobbles together previously reported ideas. To do this, we here conduct a walk-through of this manuscript’s Results section, addressing each of the test systems presented there, and providing commentary on what is and what is not new in the *spotter* methodology.

We began by simulating the transcription of lone RNAPs, since their dynamics have been the focus both of single-molecule experimental studies^39,-40^ and of earlier simulation models^4,5^ that have all made important contributions to our mechanistic understanding of transcription. In designing *spotter*’s transcription module, we sought to build on the detailed mechanistic models of transcriptional kinetics previously employed in simulations, including those that account for the sequence-dependence of transcription elongation rates^3,4^. In those previous models, position-specific dwell times are an emergent property of position-specific RNAP translocation rates (which depend on the relative nucleic-acid hybridization energies of adjacent template positions) and position-specific secondary structures adopted by nascent RNAs (which depend on the template position of the RNAPs producing those RNAs); there is no guarantee that their output matches experiment. The model of transcription used there employs pause states and models of backtracking that have been presented elsewhere, but without the inclusion of gene-position-dependent dwell-times that reflect the often rough energy landscape over which transcription appears to occur. One of the principal innovations of the present work is to show that the residue-level description of transcription that is provided by NET-seq data can be combined with single-molecule data on pausing kinetics to yield a simulation model that makes these data mutually consistent. As is described in detail in the Methods, this consistency is achieved by careful derivation of position-specific elemental pause entry rates that account for contributions to the net dwell time from longer-lived, but less frequently accessed, pause states.

We then considered how inclusion of DNA supercoiling between two transcribing RNAPs would affect their rates of transcription. *spotter* is not the first simulation model to be used to explore a connection between supercoiling and transcription. In particular, motivated by the “twin-supercoiling” model of transcription, in which positive supercoiling density (σ) accumulates downstream of a translocating RNAP and negative supercoiling density is generated upstream of it^33^, experimental studies have demonstrated that transcription is central in defining supercoiling density in different regions across the genome^56^ and may drive cooperative behavior among transcribing polymerases^16^. A number of simulation studies have explored the relationship between DNA supercoiling and transcription initiation rate^11,62–64^, and, beginning with pioneering work by Heberling et al.^9^ which built on single-molecule studies of transcription under torsion^46^, the relationship between supercoiling and transcription elongation rates^10,12–14^. To our knowledge, however, no simulation method capable of representing both RNAP backtracking and DNA supercoiling has been reported. In *spotter*, we extend previous models to build a supercoiling module that includes this coupling and that is directly integrated with a model of co-transcriptional translation. In addition, our model differs from previous models in its inclusion of a detailed system of reactions resulting in RNA chain elongation; sequence-specific rates for these reactions; an elaborate scheme for representing the activity of gyrase and topoisomerase I as processive processes; and a method for representing supercoiling-dependent rotational relaxation of RNA polymerases.

Having shown that even in the presence of supercoiling-mediated communication between RNAPs, significant potential exists for traffic to grind to a halt (Figure 6A), we used *spotter* to propose and explore a potential mechanism by which congestion could be relieved. Specifically, we invoked the possibility that the torque derived from the build-up of supercoiling between two RNAPs could assist in returning a paused RNAP to an on-pathway state, and thereby provide a mechanism by which a lagging RNAP could ‘wake up’ a leading RNAP even at a distance of ∼300 bps. The exploration of such a mechanism is only possible in *spotter* owing to its modeling of the advanced pause states observed in single-molecule studies. At this point, the notion of an assistive “release torque threshold” remains a hypothesis to be tested, or refuted, experimentally, but we think it provides a good illustration of the way in which simulation models that are sufficiently sophisticated can be used as devices for hypothesis generation. We note in passing that the release torque threshold that we assume here is entirely reasonable given that is within the range of torques generated by transcription and is a value at which the frequency and duration of shorter-lived elemental pauses are reduced in single-molecule experiments^46^.

To provide a further illustration of *spotter*’s modeling of supercoiling, we next considered the transcription of the *rpoB* gene embedded within a 10 kbp isolated topological domain. Again, previous simulation work has frequently focused on these domains as barriers to the diffusion of supercoiling that affect transcription initiation rates^62=64^. More recently, simulation studies have focused on the role of enclosing domains on transcriptional elongation rates^10–14^. Here, we build on models that have represented the activity of these enzymes either implicitly^10,13,62^ or, in various degrees of detail, explicitly^11–12,64^. The principal innovation in *spotter* with regard to modeling of topological domains is its sophisticated treatment of the topoisomerases that, *in vivo*, do much to mitigate the effects of supercoiling that necessarily accompany transcription. Specifically, we have incorporated experimentally-derived parameters characterizing the lag and burst phases that these DNA-relaxing enzymes undergo, allowing us to model accurately the sometimes-abrupt changes in supercoiling density experienced by RNAPs transcribing the template the enzymes act on.

To provide an illustration of *spotter*’s modeling of translation we considered first the monocistronic *gapA* operon. Although seminal simulation studies^26–29^ that include both transcription and translation represent an important step forward in addressing the latter question, currently-available models omit elements crucial to the coordination of prokaryotic transcription and translation. None of the models includes DNA supercoiling, backtracking, or RNAP rotational dynamics and, to render simulation systems more tractable, models have incorporated artificial reactions to automatically load ribosomes on completion of transcription of the ribosome binding site^27^, assigned transcription and translation to separate reaction compartments^26^, omitted RNAP pausing^28^, or limited simulations to one RNAP^29^. Output characterizing translation from these reduced models is also typically limited to inventories of full-length proteins.

Finally, we showed how *spotter* allows integrated models of transcription and translation to be applied to polycistronic operons, considering here the bicistronic *dusB-fis* operon and the tricistronic *marRAB* operon. Although *spotter* is not the only simulation method capable of modeling polycistronic operons^28^, previous models have relied on single-state models of transcription and translation. The software we report here is also unique in providing detailed trajectories characterizing moment-by-moment translational usage of each gene within such an operon. In its inclusion of the bidirectional interaction between transcription and DNA supercoiling, *spotter* is also particularly well-suited to modeling the role that the extensive translational cargo carried on polycistronic operons may play in determining transcriptional elongation rates.

While we have provided a number of illustrative applications of *spotter*, it is worth noting where future development will be required. One aspect that will need to be looked at is the fact that some of its input parameters are not known with any degree of certainty. Examples include topoisomerase I and gyrase binding rates, RNAP rotational relaxation rates, and gene-specific rates of accomplishing the closed-to-open transition that occurs during transcription initiation. In future work we will seek to determine which parameters are most crucial; ideally, having been identified by simulation as important to “nail down”, they might be prioritized for experimental measurement.

In addition to more narrowly defining some of the input parameters, entirely new processes are likely to require inclusion in the future. Examples include: (a) explicit modeling of RNA secondary structure: simulation models of lone RNAPs that include the effect of RNA folding on backtracking have already been reported^3,4^, but those models do not include translation and so do not account for the effect of ribosomes in their predictions of base-pairing, (b) accounting for interactions between RNA and DNA outside of the transcription bubble; although we lack clear information about the kinetics of their formation, we hope that a future iteration of the model will include these R-loops, particularly since their formation is intimately connected to DNA supercoiling^65^.

Finally, it is worth noting that while our preferred use scenario for *spotter* features systems for which next-generation sequencing data are available to allow both the transcriptional and translational landscapes to be fully described, there is no need for such data to be available in order to run simulations. For example, although the applications we have described here incorporate data only from *E. coli*, the simulation model is designed to handle analogous data characterizing any other prokaryotic organism. In fact, *spotter*’s range of applicability extends further than this, since the simulation model of *spotter* is completely agnostic regarding the level of detail in the input data, so if a user wishes to model, for example, transcription and translation occurring on completely flat landscapes (i.e. without any pauses), this is entirely possible: all that is required is that the dwell-times input by the user for each position of a gene or transcript be made identical. Similarly, if the user wishes to investigate the effects of an arbitrary sequence of pauses, this can be achieved by suitable arrangement of the input data. In other words, *spotter* provides a straightforward way of assessing the effects of entirely artificial scenarios to learn more about the coupling of transcription, supercoiling, and translation. For all these reasons, we anticipate that *spotter* will be a valuable resource for investigators interested in modeling transcription and translation as they occur *in vivo*.

## DATA AVAILABIILTY

All computer code necessary to run the simulations described here will be made available to reviewers at the time of manuscript review. Upon acceptance of the manuscript for publication, the computer code will be available to the community at the following GitHub repository (https://github.com/Elcock-Lab/spotter).

## SUPPLEMENTARY DATA

Supplementary Data are available at NAR Online.

## ACKNOWLEDGEMENT

This research was supported in part through computational resources provided by The University of Iowa, Iowa City, Iowa.

